# E3 ubiquitin ligase SYVN1 is a key positive regulator for GSDMD-mediated pyroptosis

**DOI:** 10.1101/2021.07.21.453219

**Authors:** Yuhua Shi, Yang Yang, Weilv Xu, Wei Xu, Xinyu Fu, Qian Lv, Jie Xia, Fushan Shi

## Abstract

Gasdermin D (GSDMD) participates in activation of inflammasomes and pyroptosis. Meanwhile, ubiquitination strictly regulates inflammatory responses. However, how ubiquitination regulates Gasdermin D activity is not well understood. In this study, we show that pyroptosis triggered by Gasdermin D is regulated through ubiquitination. Specifically, SYVN1, an E3 ubiquitin ligase of gasdermin D, promotes GSDMD-mediated pyroptosis. SYVN1 deficiency inhibits pyroptosis and subsequent LDH release and PI uptake. SYVN1 directly interacts with GSDMD, and mediates K27-linked polyubiquitination of GSDMD on K203 and K204 residues, promoting GSDMD-induced pyroptotic cell death. Thus, our findings revealed the essential role of SYVN1 in GSDMD-mediated pyroptosis. Overall, GSDMD ubiquitination is a potential therapeutic module for inflammatory diseases.

## Introduction

Pyroptosis is a form of programmed cell death, characterized by cell swelling, pore formation in cell membrane and cell lysis, releasing the cytoplasmic contents [1–4]. It plays an important role in host defense and inflammatory responses [5]. Recent studies show that pyroptosis is induced by proteolytic cleavage of Gasdermin D (GSDMD) by inflammatory caspases [2, 6, 7]. Canonical inflammasomes, including NLRP3 inflammasome, activate caspase-1. Meanwhile, lipopolysaccharide (LPS) activates non-canonical inflammasomes via the Caspase-11 in marine animals or Caspase-4 and -5 pathways in humans [8–12]. As the final downstream effector of inflammasomes activation, GSDMD is cleaved by inflammatory Caspases at the junction between the N-terminal cytotoxic domain (GSDMD-p30) and C-terminal autoinhibitory domain (GSDMD-p20) [5, 13, 14]. GSDMD-p30 then binds (cellular) phospholipids and oligomerizes to form 10-20 nm pores on the plasma membrane, triggering pyroptotic cell death [15–19].

Post-translational modification of inflammasome components via ubiquitin (Ub) is critical for regulation of inflammasomes activation [20, 21]. Ubiquitination of NLRP3, Caspase-1/11, ASC, IL-1β and other inflammasome components regulates several essential nodes in the regulatory networks [20–27]. E3 ubiquitin ligases such as Pellino 2 mediates K63-linked polyubiquitination and NLRP3 inflammasome activation [28]. In addition, HUWE1 stimulates inflammasomes activation by promoting K27-linked polyubiquitination of NLRP3, AIM2, and NLRC4, which strengthen host defense against bacterial infection [29]. Gasdermin B (GSDMB), a member of gasdermin family, has also been recently found to participates in ubiquitination-mediated pyroptosis that blocks bactericidal functions of NK cells [30]. Given that pyroptosis regulation is important in signaling of cell death, GSDMD ubiquitination may broadly be thought to regulate the function of GSDMD and the activities of different inflammasomes [31]. Importantly, the function and mechanisms underlying post-translational modifications of GSDMD such as ubiquitination, deubiquitination and phosphorylation are still unknown.

Synoviolin (SYVN1), also known as Hrd1, is one of the RING E3 ligases [32]. Given that SYVN1 target numerous substrates such as a pro-apoptotic factor (IRE1) [33], B lymphocyte–induced maturation protein 1 (BLIMP1) [34] and mitochondrial antiviral signaling (MAVS), it performs distinct functions in different cells [35]. Furthermore, SYVN1 regulates ER-stress-induced cell death by promoting ubiquitination and degradation of IRE1. Overall, this regulates survival and death of cells [33]. Moreover, SYVN1 promotes T-cell immunity [36] B-cell immunity [37] and regulates TLR-induced inflammation through K27-linked ubiquitination and inactivation of Usp15 [38]. However, the role of SYVN1 in pyroptosis is not well understood.

In this study, we found SYVN1 promotes canonical and non-canonical pyroptosis through GSDMD ubiquitination. Particularly, SYVN1 induces GSDMD-induced pyroptotic cell death by promoting K27-linked polyubiquitination of GSDMD on K203 and K204 residues. Our findings revealed a new mechanism of pyroptotic cell death through post-translational modification of GSDMD.

## Results

### Ubiquitination of GSDMD

Prior this study, GSDMD ubiquitination and its effect on pyroptosis had not been reported. Herein, HEK293T cells were first transfected with plasmids expressing HA-ubiquitin and Flag-GSDMD. Immunoprecipitation was performed using anti-Flag antibody, whereas western blot was performed using anti-HA or anti-Flag. We found co-expression of HA-ubiquitin and Flag-human GSDMD induced ubiquitination of human GSDMD (Fig. 1A). Further studies confirmed that both mouse and porcine GSDMD proteins can also be ubiquitinated (Fig. 1B, 1C). Further experiments revealed that endogenous human GSDMD can also be modified by ubiquitination (Fig. 1D, 1E). Interestingly, LPS/nigericin or TLR1/2 ligand Pam3CSK4 treatment followed by LPS transfection substantially increased ubiquitination of GSDMD in THP-1 cells, which activated canonical and non-canonical inflammasomes (Fig. 1D-G). Accordingly, we hypothesized that GSDMD ubiquitination is a critical and efficiently regulated process.

**Fig. 1.**
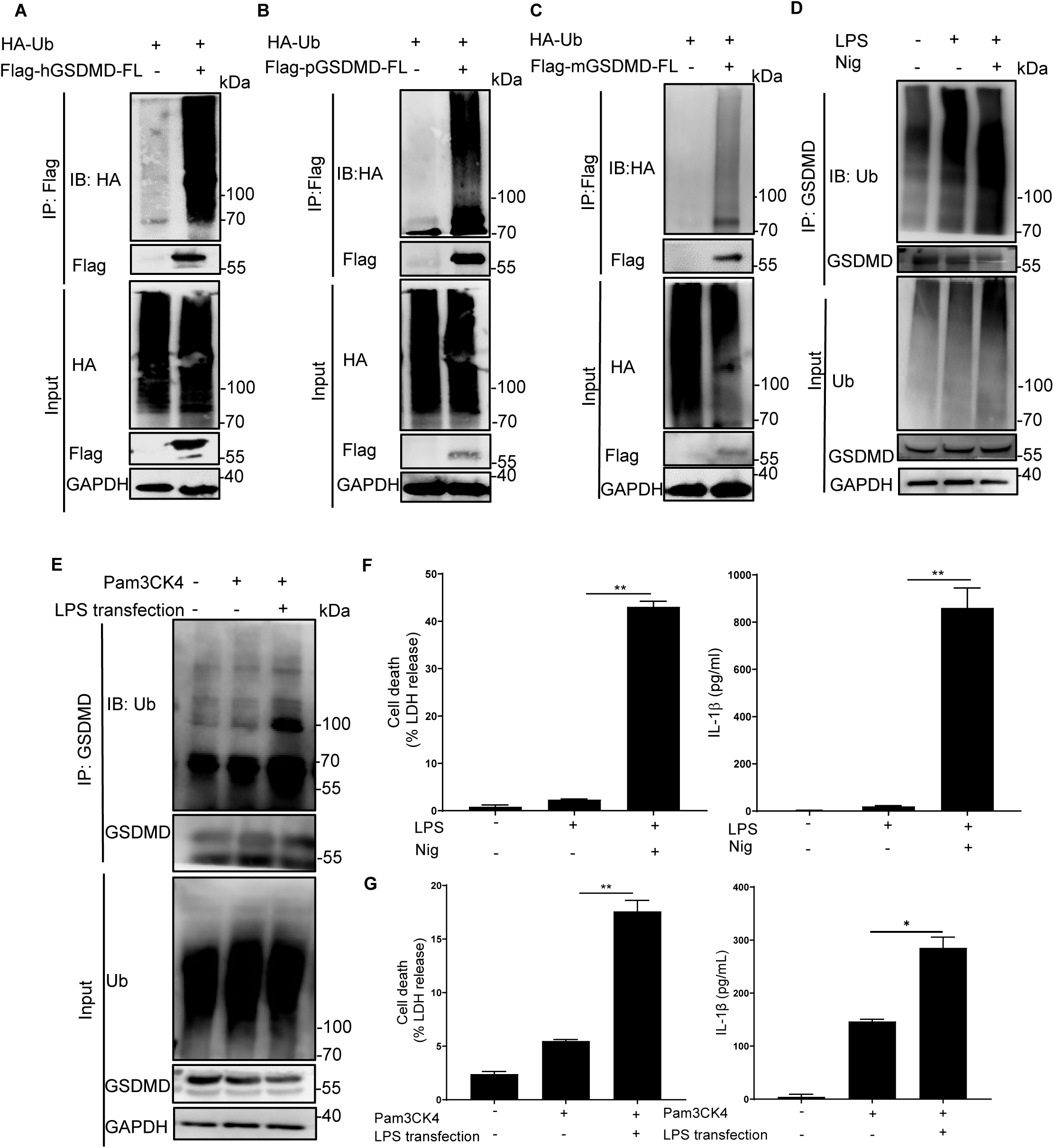
GSDMD can be modified by ubiquitination. **A,** Lysates from HEK293T cells co-transfected with HA-Ub along with Flag-human GSDMD (hGSDMD) or not, were subjected to immunoprecipitation with Flag antibody followed by western blot analysis using anti-HA antibody. **B,** Immunoprecipitation and western blot analysis of HEK293T cells co-transfected with HA-Ub and Flag-porcine GSDMD (pGSDMD) using anti-Flag and anti-HA antibodies, respectively. **C,** Immunoprecipitation and western blot analysis of HEK293T cells co-transfected with HA-Ub and Flag-mouse GSDMD (mGSDMD) using anti-Flag and anti-HA antibodies, respectively. **D**, **F**, THP-1 cells were primed with LPS (500 ng/ml) for 4 h, and then treated with or without Nigericin (10 μ Cell lysates were immunoprecipitated with anti-GSDMD antibody, followed by western blot analysis with anti-GSDMD and anti-Ub antibodies, respectively. Secretion of IL-1β was analyzed using ELISA. LDH release was analyzed using LDH release assay. **E**, **G**, THP-1 cells were incubated with Pam3CK4 (500 ng/ml) for 4 h, and then transfected with or without LPS (2 μg/ml) for 6 h. Cell lysates were immunoprecipitated with anti-GSDMD antibody, followed by western blot analysis with anti-GSDMD and anti-Ub antibodies. IL-1β secretion in THP-1 cells was analyzed using ELISA, whereas LDH release was analyzed using LDH release assay.

### Identification of SYVN1 as a GSDMD-interacting E3 ligase

Since E3 ligase plays an indispensable role in ubiquitination process, we explored the involvement of this enzyme in GSDMD ubiquitination. Analysis of UbiBrowser database revealed that SYVN1 is one of the most important E3 ligases involved in GSDMD ubiquitination (Fig. S1A). HEK293T cells were therefore transfected with Myc-tagged SYVN1 and Flag-GSDMD. We found GSDMD interacted with SYVN1 (Fig. 2A). IP LC MS/MS analysis revealed comparable findings (Fig. 2D, S1B). Further analyses revealed that GSDMD also interacted with SYVN1 in THP-1 cells (Fig. 2B, 2C). Indirect immunofluorescence further revealed that interaction between SYVN1 and human GSDMD occurred in the cytoplasm (Fig. 2E). In general, interaction between SYVN1 and GSDMD is both endogenous and exogenous.

**Fig. 2.**
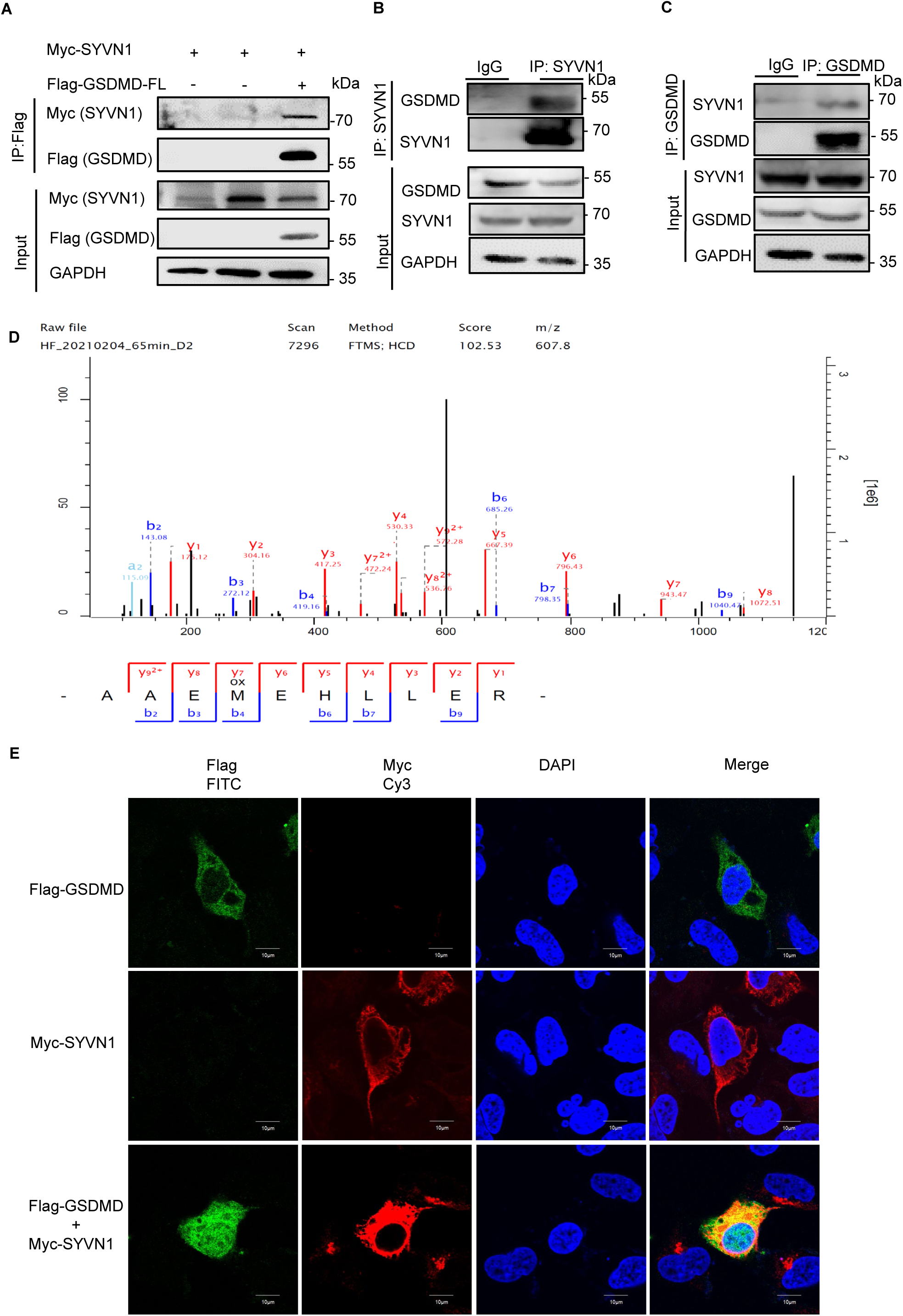
SYVN1 interacts with endogenous and exogenous GSDMD. **A,** HEK293T cells co-transfected with pcDNA3.1-SYVN1-Myc and p3xFlag-GSDMD were lysed and immunoprecipitated with anti-Flag antibody. Both the immunoprecipitates (IP) and whole cell lysates (Input) were subjected to gel electrophoresis and analyzed by western blot with anti-Myc and anti-Flag antibodies, respectively. **B-C,** Immunoprecipitation analysis of THP-1 cells using anti-SYVN1 or anti-GSDMD antibodies. Western blot of both IP and total proteins in THP-1 cells was performed using anti-SYVN1 and anti-GSDMD antibodies, respectively. **D,** HEK293T cells co-transfected with pcDNA3.1-SYVN1-Myc and p3xFlag-GSDMD were lysed and immunoprecipitated with anti-Flag antibody. The potential GSDMD-binding proteins in HEK293T cells were evaluated using Co-IP and MS analysis. The MS/MS spectrum of 71-AAEMEHLLER-80 is shown. Observed b- and y-ion series are indicated. **E,** Immunostaining analysis of HEK293T cells after transfection with pcDNA3.1-SYVN1-Myc and p3xFlag-GSDMD using anti-Flag and anti-Myc antibodies. Subcellular localizations of Flag-GSDMD (green), Myc-SYVN1 (red), and DAPI (blue), a nucleus marker, were observed using a confocal microscopy. Scale: 1 bar represents 10 μm.

### Reconstruction of canonical and non-canonical pyroptosis *in vitro*

Given that GSDMD is the key effector of pyroptosis downstream of activated canonical and non-canonical inflammasomes [5, 6], we evaluated the role of GSDMD ubiquitination on pyroptosis. Since HEK293T cells do not express endogenous GSDMD [39, 40], we reconstructed canonical and non-canonical pyroptosis in HEK293T cells. Expression of both GSDMD and Caspase1/4 significantly induced LDH production (Fig. 3A, 3B). Western blot revealed that Caspase-1 and Caspase-4 cleaved the full-length GSDMD into active N-terminal GSDMD-p30 domain (Fig. 3A, 3B) and obvious propidium iodide (PI) staining was observed by fluorescence microscopy (Fig. 3C, 3D). Same results were observed in cell death and PI staining analyses of GSDMD-p30 triggered pyroptosis (Fig. 3A-D). However, the expression of GSDMD-p30 was not detectable, in agreement with the hypothesis that the protein might have a toxic effect on the host cell [41]. Overall, these results show that the canonical and non-canonical pyroptosis were successfully reconstructed *in vitro*.

**Fig. 3.**
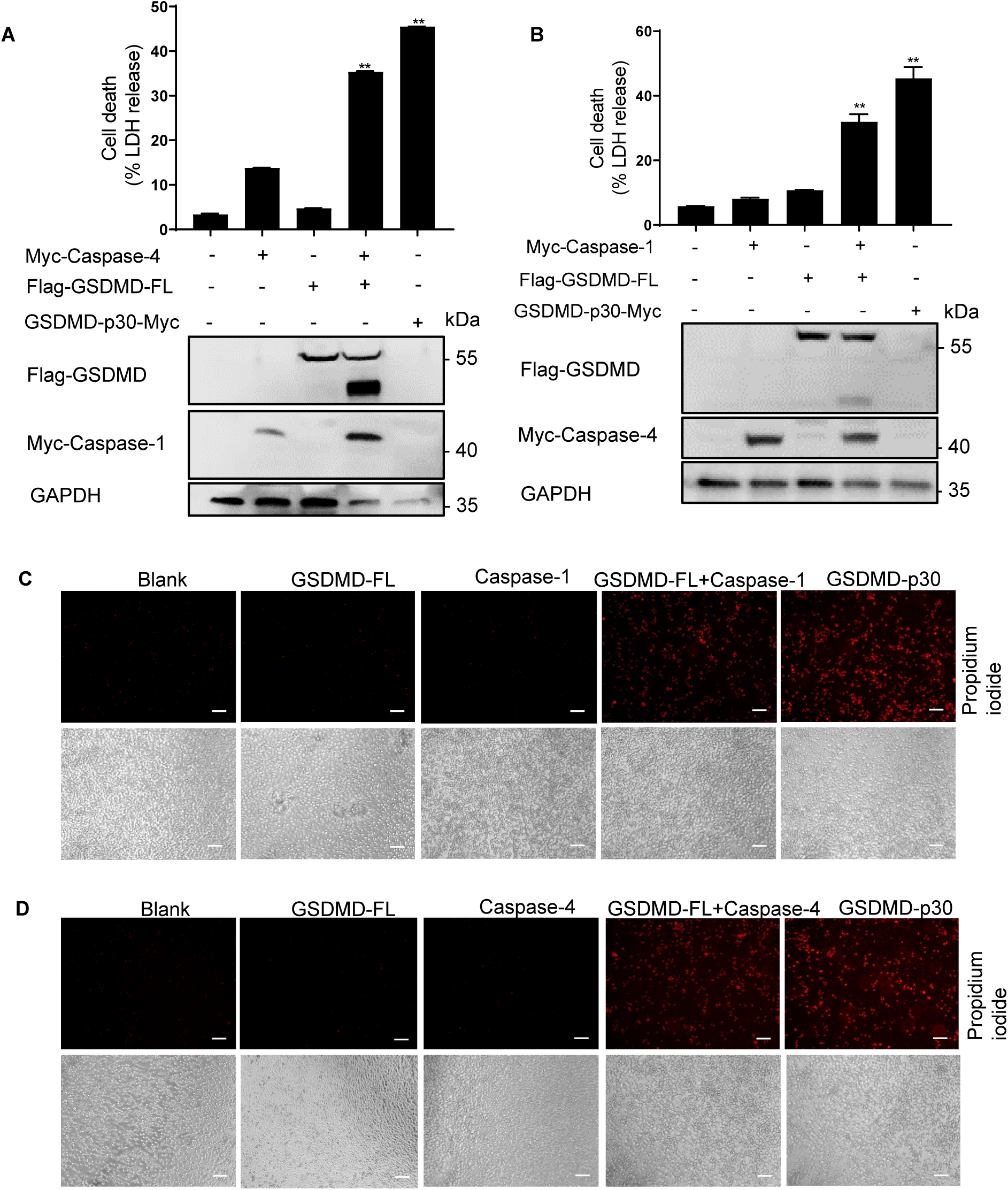
Reconstruction of canonical and non-canonical pyroptosis *in vitro*. **A,** HEK293T cells were co-transfected with pCMV-Myc-Caspase-1 (600 ng) and p3XFlag-hGSDMD-FL (600 ng). pcDNA3.1-hGSDMD-p30-Myc (600 ng) was used as a positive control. Supernatants were analyzed using LDH assay. Cell lysates were normalized for proteins contents and analyzed using western blot using antibodies specific for Flag, Myc and GAPDH. **B,** HEK293T cells were co-transfected with pCMV-Myc-Caspase-4 and p3XFlag-hGSDMD-FL. The pcDNA3.1-hGSDMD-p30-Myc was used as positive controls. Anti-Flag, Myc and GAPDH antibodies were used for western blot analyses. **C-D,** Pyroptosis of HEK293T cells after GSDMD cleavage as observed under bright-field and epifluorescent microscopy. Scale: 1 bar represents 100 μm.

### SYVN1 promotes GSDMD-mediated pyroptosis

To determine whether SYVN1 regulates GSDMD-mediated pyroptosis, we evaluated LDH release and PI staining (red) based on the reconstruction of canonical and non-canonical pyroptosis. SYVN1 overexpression markedly increased canonical and non-canonical pyroptosis of cells under expressing Caspase-1/4 and GSDMD (Fig. 4A, 4D and S2A). To confirm that SYVN1 performed its function independent of Caspase-1/4, Caspase1/4 was transfected into HEK293T cells 12 h after GSDMD and SYVN1 co-transfection. Pyroptosis was assessed based on LDH release and PI staining (red). LDH release and PI staining were significantly increased compared with the group without SYVN1 after 6 h, 18 h and 24 h of transfection (Fig. 4B, S2B-2C). In addition, SYVN1 interacted with GSDMD but not with Caspase1/4 (Fig. 4H). Accordingly, we speculated that SYVN1 promotes pyroptosis by directly targeting GSDMD. To validate this hypothesis, we evaluated whether direct SYVN1 targeting of GSDMD N-terminal (GSDMD-p30) induces pyroptosis. Of note, transfection of SYVN1 with GSDMD-p30 markedly enhanced LDH release and PI uptake compared with the group without SYVN1 transfection (Fig. 4C, 4D). These results demonstrated that SYVN1 accelerates pyroptosis mainly by targeting the GSDMD-p30 fragment. To further investigate whether E3 ubiquitin ligase activity of SYVN1 is responsible for regulating GSDMD-induced pyroptosis, we assessed the effects of SYVN1 and SYVN1-C329S on pyroptosis in HEK293T cells. Based on previous reports, the cysteine residue at position 329 is critical, and its mutation to serine (C329S) abolishes SYVN1 E3 activity [38, 42]. We found SYVN1-C329S which lacks E3 ligase activity could not increase LDH release (Fig. 4E-G). To determine whether GSDMD ubiquitination activates pyroptosis, HEK293T cells were co-transfected with GSDMD-p30 and wild-type Ub. It was found that overexpression of Ub significantly promoted GSDMD-p30-mediated pyroptosis (Fig. S2D, S2E), indicating that ubiquitination of GSDMD facilitates pyroptosis. Importantly, the addition of Ub further promoted SYVN1 enhanced GSDMD-p30-mediated pyroptosis (Fig. 4I). Collectively, these findings demonstrated that SYVN1 promotes pyroptosis through ubiquitination of GSDMD.

**Fig. 4.**
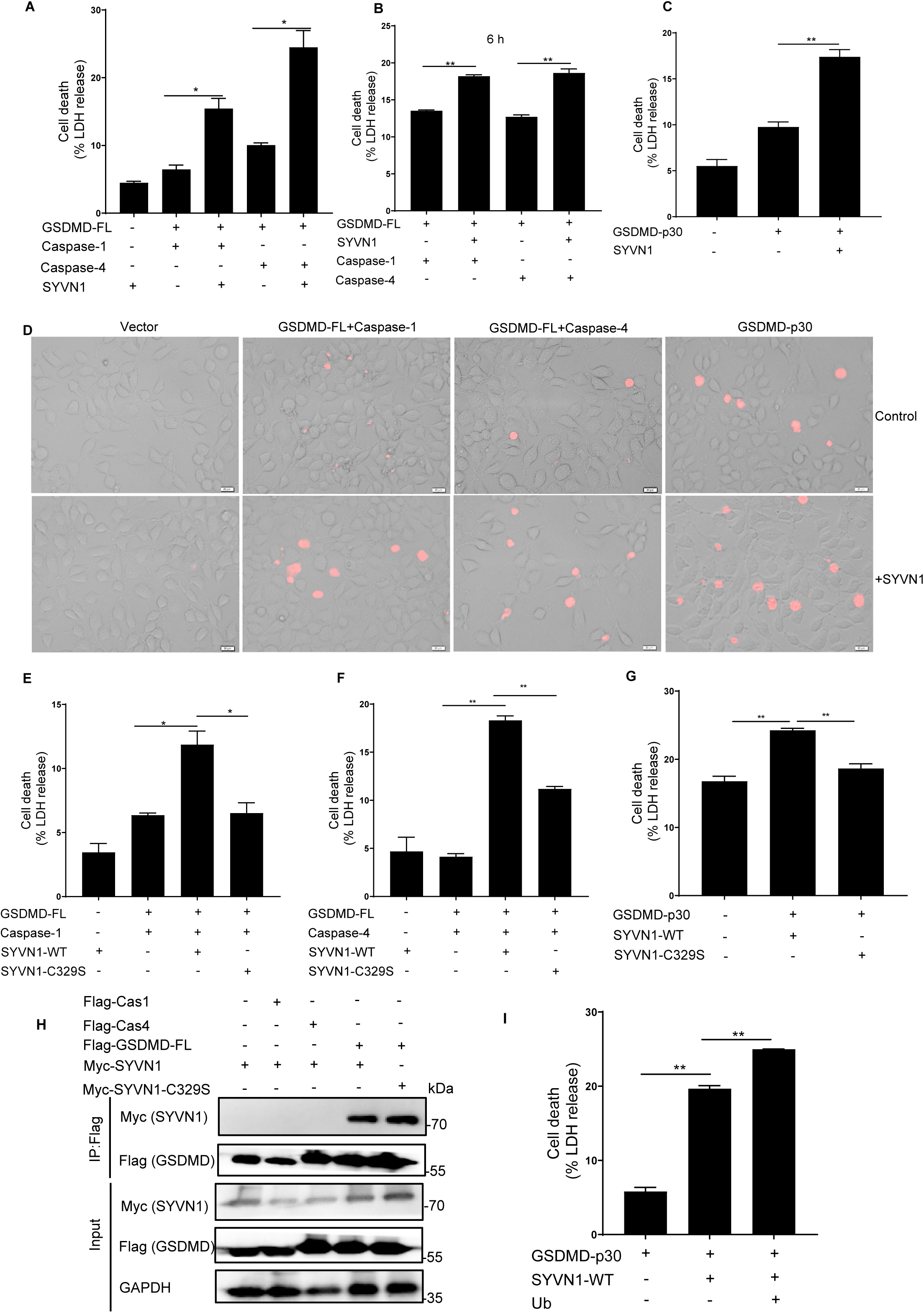
Overexpression of SYVN1 promotes GSDMD-triggered pyroptosis. **A, D,** HEK293T cells were transfected with pCMV-Myc-Caspase-1/4 (100 ng), p3XFlag-hGSDMD-FL (200 ng) and p3XFlag-SYVN1 (600 ng) or control vector. The supernatants were collected and analyzed by LDH release assay and cell staining with PI after 24 h transfection. PI analysis was performed using 2.5 μg/ml. Scale: 1 bar represents 20 μm. **B,** HEK293T cells were transfected with Myc-SYVN1 (600 ng) and p3XFlag-hGSDMD-FL (200 ng) for 12 h, and then transfected with Caspase-1/4 (100 ng) for 6 h. **C-D,** HEK293T cells were transfected with Myc-SYVN1 (600 ng) and hGSDMD-p30-Myc (100 ng). Supernatants were collected and analyzed by LDH release assay and cell staining with PI at 24 h after transfection. **E-F,** LDH release assay of HEK293T cells after co-transfection with Caspase-1/4 (100 ng), hGSDMD-FL (200 ng) and SYVN1-WT (600 ng) or SYVN1-C329S (600 ng). **G,** LDH release assay of HEK293T cells co-transfected with hGSDMD-p30 (100 ng) and SYVN1-WT (600 ng) or SYVN1-C329S (600 ng). The supernatants were collected and analyzed by LDH release assay at 24 h after transfection. **H,** Immunoprecipitation (IP) and immunoblot (IB) analysis of HEK293T cells were co-transfected with pcDNA3.1-SYVN1-Myc and p3XFlag-hGSDMD-FL or p3XFlag-Caspase-1/4. IP was performed using anti-Flag antibody, whereas IB was performed using anti-Myc antibody. **I,** LDH release of HEK293T cells co-transfected with hGSDMD-p30, SYVN1 and Ub. The analysis was performed at 24 h after transfection.

### Inhibition of SYVN1 impairs GSDMD-mediated pyroptosis

To further confirm the role of SYVN1 on pyroptosis, we designed three siRNAs targeting SYVN1. The efficiency of SYVN1 knock-down was shown in Fig. 5A, and we ultimately selected siSYVN1 #3 for further studies because of its better knockdown efficacy (Fig. 5A). LDH assay and PI uptake revealed that SYVN1 knockdown inhibited the effect of GSDMD-p30 on pyroptosis in HEK293T cells (Fig. 5B, 5C, S3) To further investigate the role of SYVN1 in pyroptosis, we generated SYVN1 knockout HEK293T cells by CRISPR/Cas9-mediated genome editing (according to reference [43]). The efficiency of SYVN1 knock-out was shown in Fig. 5D, and we finally selected sgSYVN1 #2 for further research because of its better knockout effect. Consistent with the results of knockdown of SYVN1, knockout of SYVN1 in HEK293T cells significantly suppressed canonical, non-canonical pyroptosis and GSDMD-p30 activation (Fig. 5E). Collectively, these findings further demonstrated that SYVN1 promotes GSDMD-mediated pyroptosis.

**Fig. 5.**
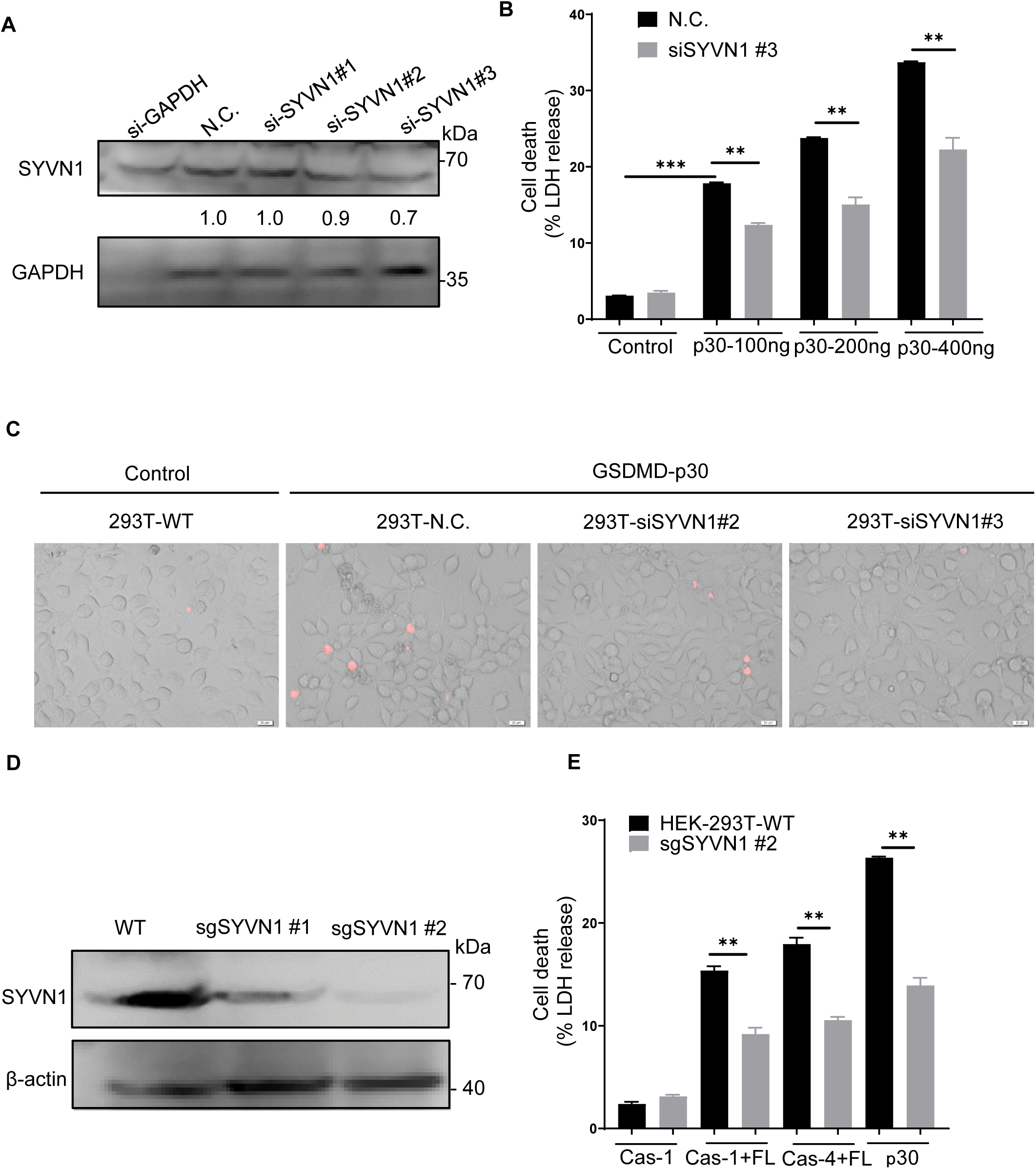
Effect of inhibiting SYVN1 on GSDMD-mediated pyroptosis. **A,** Western blot analysis for efficiency of SYVN1 expression inhibition in HEK293T cells using siSYVN1. The analysis was performed using anti-SYVN1 antibody. **B-C,** HEK293T cells were transfected siSYVN1-control (50 nM) or siSYVN1 (50 nM), 48 h after transfection, cells were transfected with hGSDMD-p30. The supernatants were collected and analyzed by LDH release assay. After 24 h transfection, the cells were staining with PI (2.5 μg/ml) and analyzed by Fluorescence microscopy. **D,** The efficiency of sg-SYVN1 was analyzed by western blot with anti-SYVN1 antibody in HEK293T cells obtained from WT and SYVN1-KO HEK293T cells. **E,** HEK293T WT and SYVN1-KO cells were transfected with Caspase-1/4 and hGSDMD-FL or hGSDMD-p30 alone. The supernatants were collected and analyzed by LDH release assay.

### SYVN1 ubiquitinates GSDMD with K27-linked polyubiquitin chains

Considering that SYVN1 regulate pyroptosis in an E3 ligase activity-dependent manner, we speculated SYVN1 mediates polyubiquitination of GSDMD. As such, we co-expressed Myc-SYVN1-WT or the E3 ligase-dead mutant SYVN1 C329S with HA-ubiquitin (HA-Ub) and Flag-GSDMD in HEK293T cells. We found compared to C329S mutant, overexpression of SYVN1-WT substantially increased GSDMD ubiquitination (Fig. 6A), suggesting that E3 ubiquitin ligase activity is necessary for SYVN1 ubiquitination of GSDMD. Since K48 and K63 are the most studied canonical ubiquitination, we further investigated the role of these two polyubiquitination chains in ubiquitination of GSDMD. HEK293T cells were transfected with Flag-GSDMD and HA-K48-ubiquitin, HA-K63-ubquitin, HA-K48R-ubiquitin or HA-K63R-ubquitin. It was found SYVN1 had no effect on K48 and K63-mediated polyubiquitination of GSDMD (Fig 6B). Also, SYVN1 promoted K48R and K63R-mediated polyubiquitination, suggesting that SYVN1 induced addition of other ubiquitination chains on GSDMD. Given that seven lysine residues (K6, K11, K27, K29, K33, K48 and K63) have been reported to form polyubiquitin chains [44–48], we sequentially co-transfected these HA-tagged ubiquitin mutants (with only one of the seven lysine residues retained as lysine, and the other six replaced with arginine) with Flag-tagged GSDMD with or without SYVN1 into HEK293T cells. Co-immunoprecipitation assay revealed that K27 ubiquitin mutant markedly increased polyubiquitination of GSDMD (Fig. 6C). However, overexpression of SYVN1 had no effect on GSDMD polyubiquitination in K27R mutant transfected cells (Fig. 6D). These findings suggest that SYVN1 promoted GSDMD-mediated pyroptosis may through K27-linked polyubiquitination of GSDMD. Indeed, overexpression of Ub-K27 promoted pyroptosis induced by Caspase-1/4 mediated GSDMD cleavage and GSDMD-p30 activation (Fig. 6E-F). Thus, SYVN1 mediates K27-linked polyubiquitination of GSDMD.

**Fig. 6.**
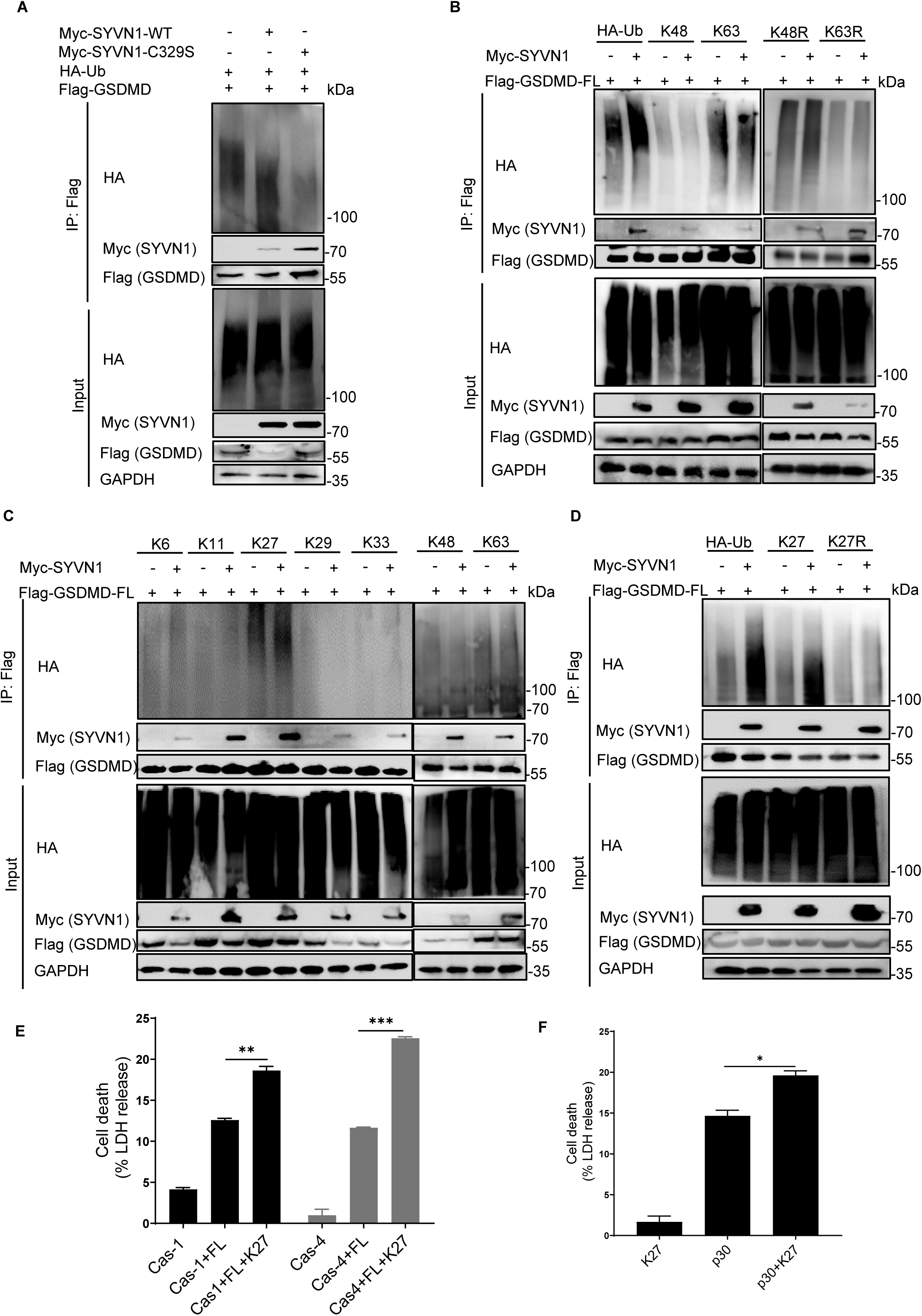
SYVN1 ubiquitinates GSDMD with K27-linked polyubiquitin chains. **A,** Immunoprecipitation analysis in HEK293T cells co-expressing Flag-GSDMD and HA-Ub together with Myc-SYVN1 or Myc-SYVN1-C329S. Anti-Flag immunoprecipitates were analyzed using western blot with indicated antibodies. The expression levels of the transfected proteins were analyzed using western blot with indicated antibodies. **B,** Immunoprecipitation analysis of HEK293T cells co-expressing Flag-GSDMD and Myc-SYVN1 together with HA-Ub (K48, K63, K48R or K63R) as indicated. Western blot analysis was perfomed using anti-Myc or anti-Flag antibodies. Expression of the transfected proteins was analyzed using western blot, using corresponding antibodies. **C,** Immunoprecipitation analysis of HEK293T cells expressing Flag-GSDMD and Myc-SYVN1 together with HA-Ub (K6, K11, K27, K29, K33, K48, or K63 only) as indicated. Anti-Flag immunoprecipitates were analyzed using western blot with anti-Myc or anti-Flag antibodies. Expression of the transfected proteins was analyzed using western blot, using corresponding antibodies. **D,** Immunoprecipitation analysis of HEK293T cells expressing Flag-GSDMD and Myc-SYVN1 together with HA-Ub (K27 or K27R) as indicated. Anti-Flag immunoprecipitates were analyzed using western blot with indicated antibodies. Expression of the transfected proteins was analyzed using western blot, using corresponding antibodies. **E,** LDH release assay of HEK293T cells transfected with Caspase-1/4, GSDMD and Ub-K27 or control vector. **F,** LDH release assay of HEK293T cells transfected with hGSDMD-p30 and Ub-K27 or control vector.

### K203 and K204 residues of GSDMD are polyubiquitinated by SYVN1

To determine sites of GSDMD polyubiquitination by SYVN1, ubiquitination in HEK293T cells was induced through expression of Flag-GSDMD, SYVN1-Myc, and HA-ubiquitin. Controls expressed SYVN1-C329S-Myc, Flag-GSDMD and HA-ubiquitin. After immunoprecipitating GSDMD with anti-Flag purification beads, the protein was separated using SDS-PAGE and visualized after Coomassie blue staining (Fig. 7A). To analyze ubiquitinated GSDMD sites by mass spectrometry (MS), the gel regions (Fig. 7A) which might contain ubiquitin-modified proteins were excised, digested with trypsin, and analyzed by liquid chromatography-MS. A database search of the MS/MS spectra revealed that GSDMD is the predominant protein identified in the samples. Representative spectra demonstrated that the K51 and K299 residues of GSDMD in gel were modified with ubiquitin (Fig. S4A). We also used the UbPred program (http://www.ubpred.org) [49–51], a predictor of potential ubiquitin binding sites in proteins to predict the potential ubiquitination sites of GSDMD. This analysis revealed other five putative target lysine residues: K103, K203, K204, K235 and K248 (Fig. S4B). We then generated Flag-GSDMD K51R, K103R, K203R, K204R, K235R, K248R and K299R mutants in which each of these lysine residues was replaced with arginine (R). To determine if these potential ubiquitination sites altered pyroptotic function, HEK293T cells were co-transfected with wild-type GSDMD or Flag-GSDMD mutants and Caspase1/4. LDH release and pore formation assay revealed that pyroptosis was significantly low in GSDMD-K103R, GSDMD-K203R, GSDMD-K204R, GSDMD-K235R and GSDMD-K248R mutants, but not in GSDMD-K51R and K299R mutants (Figs. 7B, 7C, S5A). GSDMD-D275A, which is the only cleavage site of Caspase1/4, was used as the positive control [5, 13]. As expected, Caspase-1/4 did not cleave GSDMD-D275A, and the mutant (GSDMD-D275A) had no effect on pyroptosis (Figs. 7B, 7C). Moreover, mutations in GSDMD-p30 ubiquitination sites substantially decreased GSDMD-p30-mediated pyroptosis (Fig. 7D , S5B). GSDMD-p30-C191A was used as a positive control, given that mutation (C191A) in this amino acid decreases oligomerization and inhibits pyroptosis [40, 52, 53]. Our results suggest that amino acids GSDMD-K103, GSDMD-K203, GSDMD-K204, GSDMD-K235 and GSDMD-K248 are critical for pyroptosis.

**Fig. 7.**
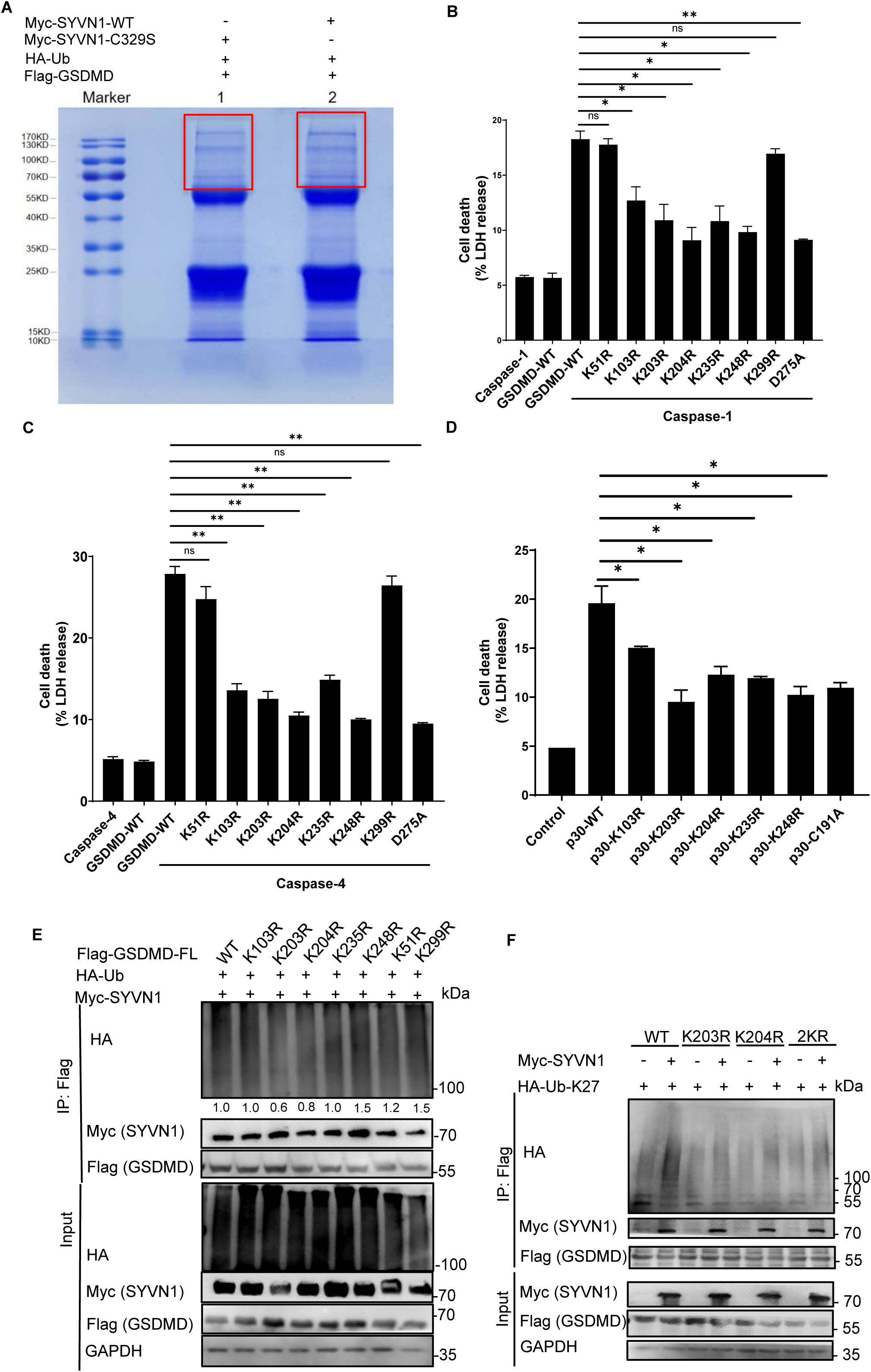
SYVN1 polyubiquitinates K203 and K204 residues of GSDMD . **A,** Coomassie blue-stained bands of polyubiquitinated GSDMD. HEK293T cells were co-transfected with Flag-GSDMD, Myc-SYVN1 or Myc-SYVN1-C329S and HA-ubiquitin (shown above the lanes). The square areas indicate regions from which proteins were removed for MS. **B-C,** LDH assay of HEK293T cells transfected with pCMV-Myc-Caspase-1/4 (300 ng), p3XFlag-hGSDMD-FL (600 ng) or GSDMD mutant (600 ng). The analysis was performed at 24 h after transfection. **D,** LDH release assay of HEK293T cells transfected with GSDMD-p30 (300 ng) or GSDMD-p30 mutants (300 ng). The analysis was performed at 24 h after transfection. **E,** Immunoprecipitation analysis of HEK293T cells expressing Flag-GSDMD or Flag-GSDMD mutants together with HA-Ub and Myc-SYVN1. Expression of proteins of interest was analyzed using western blot, using corresponding antibodies. Densitometry of the blots was measured using Image J. **F,** Immunoprecipitation analysis of HEK293T cells expressing Flag-GSDMD, K203R, K204R or 2KR mutants along with HA-Ub-K27 and Myc-SYVN1. Western blot was performed using corresponding antibodies.

To further explore SYVN1-mediated ubiquitination sites on GSDMD, HEK293T cells were transfected with Myc-tagged SYVN1, Flag-tagged GSDMD or GSDMD K/R mutants and HA-tagged ubiquitin. Compared with the WT GSDMD (Fig. 7E, lane 1), SYVN1 had no effect on ubiquitination of GSDMD-K51R or K299R mutant (Fig. 7E, lane 7, 8). In contrast, even though LDH decreased in GSDMD K103R, K235R and K248R mutants (Fig. 7B, 7C), SYVN1-mediated ubiquitination of GSDMD occurred in them (Fig. 7E, lane 2, 5, 6). Specifically, Co-IP assay revealed that SYVN1-mediated GSDMD ubiquitination was partially blocked in K203R and K204R mutants (Fig. 7E, lane 3, 4). K203R/K204R mutants were then generated to further confirm K203R and K204R lysine residues are the major ubiquitination sites in GSDMD. We found arginine mutations in these two residues (2KR) did not completely prevent GSDMD ubiquitination (Fig. 7F), suggesting that SYVN1 also modifies other lysine residues. Overall, the above findings show that SYVN1 mainly ubiquitinates GSDMD at K203 and K204 residues, but there are also other lysine residues in GSDMD that are ubiquitinated by SYVN1.

## Discussion

Pyroptosis, mediated by the inflammasomes, Caspase-1 and Caspase-4/5/11, participates in the clearance of intracellular pathogens [6, 54]. Recent studies have identified Gasdermin D (GSDMD) as the primary executioner of pyroptosis [5, 6]. However, ubiquitination-mediated regulation of GSDMD activity is poorly understood. In this study, we demonstrated that ubiquitination of GSDMD plays a critical role in pyroptosis. Moreover, SYVN1 is an important E3 ubiquitin ligase that positively regulates canonical and non-canonical pyroptosis through K27-linked polyubiquitination of K203 and K204 lysine residues on GSDMD (Fig. 8).

**Fig. 8.**
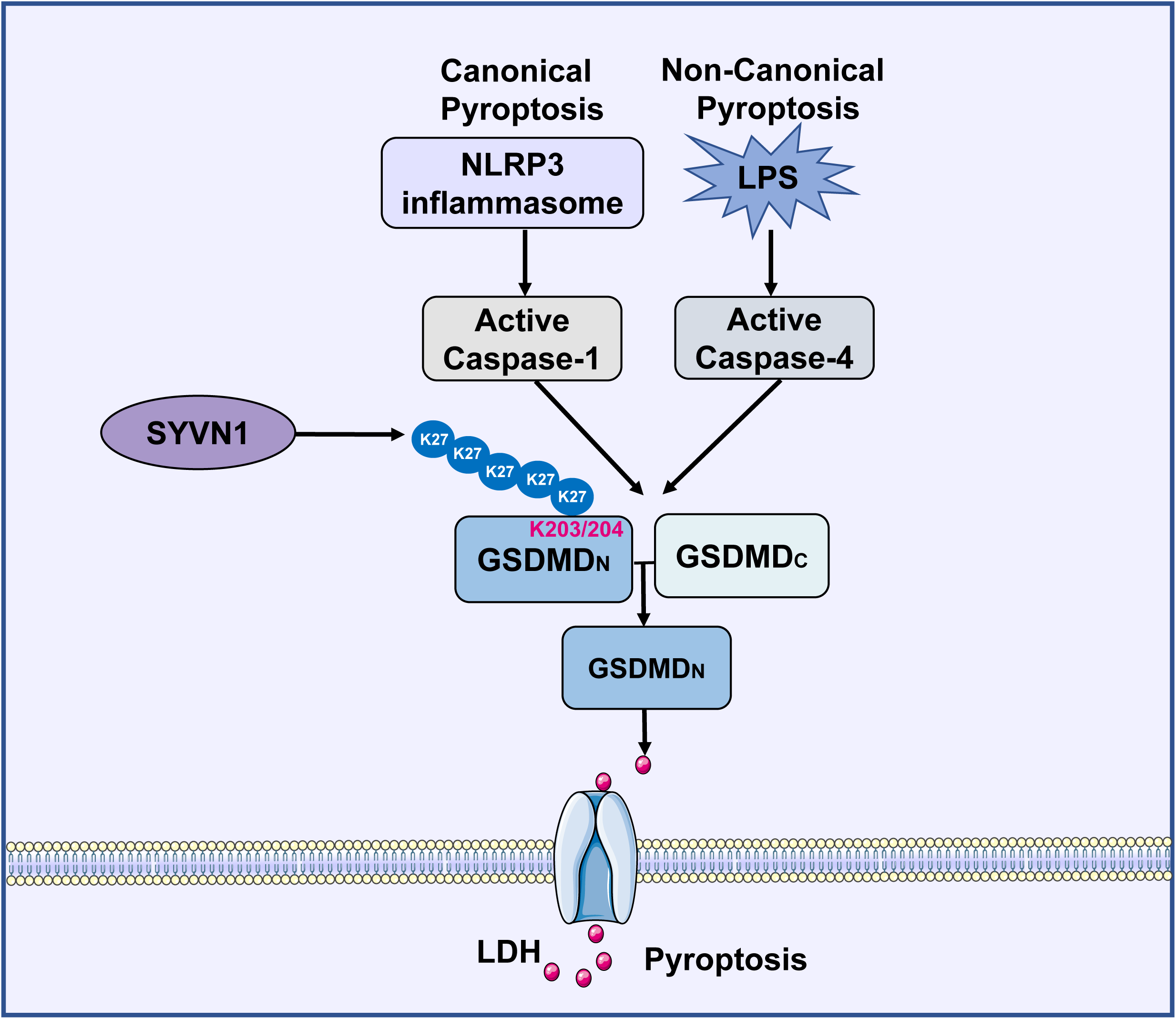
Diagram of SYVN1 regulates GSDMD-mediated pyroptosis.

Polyubiquitination is a post-translational modification process that plays critical roles in programmed cell death or NLRP3-mediated inflammasome activation through covalent linkage of ubiquitin to lysine residues in target proteins [20, 22, 55, 56]. Recent studies have found that GSDMD is essential in pyroptosis. Thus, it is reasonable to hypothesize that polyubiquitination of common components of canonical and non-canonical pyroptosis effectively regulates pyroptosis in response to infection and danger signals. Herein, we demonstrated ubiquitination of GSDMD facilitates GSDMD-mediated pyroptosis.

SYVN1 is over-expressed in synoviocytes of patients with rheumatoid arthritis, exacerbating the pathogenesis of the disease [57]. Moreover, SYVN1 regulates the functioning of immune cells, including T cells [36], B cells [37], and macrophages [38]. However, how SYVN1 regulates pyroptosis had not been previously described. We found overexpression of SYVN1 increases LDH release and pyroptosis (Fig. 4), and as such, loss of SYVN1 function impairs GSDMD-mediated pyroptosis (Fig. 5). In general, SYVN1 promotes GSDMD-mediated pyroptosis.

There were multiple K residues in GSDMD that can be ubiquitinated based on the prediction by Ubipred online software. These residues could be ubiquitinated by SYVN1 and other E3 ubiquitin ligase enzymes, and additional studies to identify these ubiquitin ligases are needed. Herein, we found SYVN1 promotes pyroptosis mainly by interacting and ubiquitinating K203 and K204 residues in GSDMD. Mass spectrometric analysis further revealed two high-confidence ubiquitin modified residues in the N- (K51) and C-terminal (K299) domains of GSDMD (Fig. S4A). However, mutation in these residues (K51R/K299R) had no significant effect on GSDMD ubiquitination, suggesting that SYVN1 also modifies other lysine residues. Furthermore, a recent study revealed that pyroptotic cell death is independent of GSDMD-K204T modification [58]. In contrast, we found ubiquitination of GSDMD-K204R mutant inhibited GSDMD-induced pyroptosis. The discrepancy may have resulted from the presence of threonine residue, which could also be ubiquitinated [59]. Further studies are needed to explore how mutations in GSDMD ubiquitination sites influence pyroptosis *in vivo*.

Taken together, findings of this study elucidated the unique role of GSDMD polyubiquitination in pyroptosis, in particular, mediated by SYVN1. First, we found GSDMD ubiquitination occurs in several species including human, mouse, and swine. Second, we found SYVN1, an E3 ubiquitin ligase, is the primary regulator of GSDMD activity, and performs its function by promoting ubiquitination of GSDMD. Specifically, SYVN1 directly interacts with GSDMD, and mediates K27-linked polyubiquitination of GSDMD on K203 and K204 residues, promoting GSDMD-induced pyroptotic cell death. Summed together, we report a new mechanism of the growing list ubiquitin-mediated regulation, a noble mechanism of PRR signaling pathways.

## Materials and Methods

### Reagents

Anti-Flag M2 (F1804), anti-Flag M2 Magnetic Beads, anti-Myc (M5546) and anti-LPS (O11:B4) were purchased from Sigma-Aldrich. Anti-GSDMD (sc-393581) and anti-GAPDH (sc-47724) were from Santa Cruz Biotechnology. Anti-Ub (sc-8017), anti-HA (3724), anti-β-actin (3700) and the secondary antibodies used for western blot and immunofluorescence assay were purchased from Cell Signaling Technology. Anti-SYVN1 (ab170901) was purchased from Abcam, whereas Nigericin, Pam3CSK4 (tlrl-pms) were purchased from InvivoGen. ELISA kits for analysis of human IL-1β were purchased from MultiSciences. CytoTox 96 LDH-release assay kit (G1780) was purchased from Promega.

### Cell culture and stimulation

Human HEK293T cells were cultured in Dulbecco’s modified Eagle’s medium (DMEM) in a humidified incubator at 37□ with 5 % CO_2_, supplemented with 10 % FBS (Corille, C1015-05), 100 μg/ml streptomycin and 100 U/ml penicillin (Hyclone, 347 SV30010). Human monocyte cell line THP-1 was cultured in PRIM 1640 medium supplemented with 10 % FBS, 100 μg/ml streptomycin and 100 U/ml penicillin. To activate canonical inflammasomes, 5 × 10^5^ cells were plated overnight in 24-well plates, and primed for 4 h with 500 ng/ml LPS. The cells were then stimulated for 1 h using 10 μM Nigericin. For non-canonical inflammasome activation, cells were primed for 4 h with 500 ng/ml Pam3CSK4, after which the medium was replaced and cells were transfected with 2 μg/ml LPS using Lipofectamine 2000 for 6 h.

### Plasmid construction and transfection

cDNAs for transfection in plasmids were obtained by reverse transcription of total RNA from HEK293T and THP-1 cells. The cDNAs were amplified using specific primers before sub-cloning the PCR products into p3Flag-CMV-7.1, pCMV-Myc and pCDNA-3.1 vectors. HA-tagged ubiquitin was provided by Dr. Yimin Yang (Zhejiang University). The primers used in this study are shown in Supplementary Table 1.

### Induction of pyroptosis in HEK293T cells

HEK293T cells grown in 24-well plates to 80% confluence were transfected with 600 ng of p3XFlag-GSDMD and 600 ng of pCMV-Caspase1/4 using VigoFect. The supernatants were collected 24 h after transfection and analyzed for IL-1β level using ELISA.

### ELISA

Supernatants from transfected cells were assayed for human IL-1β in accordance with the manufacturer’s instructions. Each experiment was performed independently at least three times. Sample preparation for ELISA was according to the method [60].

### Cytotoxicity assay and propidium iodide staining

Culture medium of HEK293T cells co-transfected for 24 h with p3Flag-CMV-7.1, pCMV-Myc, and pCDNA-3.1 was collected and assayed for LDH release using the Cytotoxicity Detection Kit (Promega) according to the manufacturer’s manual. The HEK293T cells were cultured for 15 min with 2.5 μg/ml Propidium iodide (BD Bioscience, 556463) and observed under a fluorescence microscopy. The experiments were performed independently in triplicate.

### Western blot analysis

Cells were lysed using lysis buffer (Beyotime, P0013) supplemented with Phenylmethanesulfonyl fluoride (PMSF) (Beyotime, ST506). Total proteins in the cells were extracted and thereafter separated using 12 % SDS-PAGE. The proteins were then transferred onto PVDF membrane, hybridized with primary antibodies and probed with HRP-conjugated secondary antibodies. Proteins were detected using the image system (Clinx Science Instruments, China).

### Co-immunoprecipitation assay

Cells were lysed for 30 min in ice cold lysis buffer and centrifuged at 10,000 × g at 4°C. The supernatants were then incubated with anti-Flag binding beads (Sigma, M8823) at 4 °C. The beads were then washed for 3-5 times with cold TBS. Immune complexes were denatured for 10 min at 100 °C in 1 × SDS-PAGE loading buffer before western blot analysis.

### Confocal immunofluorescence assay

HEK293T cells were seeded in 24-well at the rate of 1 × 10^5^ per well and transfected for 24 h with p3Flag-CMV-7.1, pCMV-Myc, and pCDNA-3.1 plasmids. The cells were then fixed for 30 minutes at room temperature with 4% paraformaldehyde (Beyotime, P0098) before incubation with primary antibodies in DAPI (Beyotime, C1002). The cells were then observed using a laser scanning microscope (Olympus, IX81-FV1000).

### SYVN1 knockdown using small interfering RNA

Short interfering RNAs (genepharma) specific for human SYVN1 were transfected into THP-1 and HEK293T cells using the Lipofectamine lipo8000 reagent (Beyotime, C0533FT) according to the manufacturer’s instructions. The sequences of the siRNAs used in this study are shown in Supplementary Table 2.

### Construction of SYVN1 knockout HEK293T using CRISPR/Cas9-technique

SYVN1 knockout HEK293T cells were generated using CRISPR/Cas9 technique. Vectors expressing gRNA targeting human SYVN1 were transfected into HEK293T cells using manufacturer’s protocol. In general, based on flow-cytometric analysis of GFP levels, SYVN1 knockout was achieved in 30-50% of HEK293T cells. Single cell sorting of transfected cells was performed using flow cytometry (MOFLO XDP). gRNA sequences are shown in Supplementary Table 3.

### LC-MS/MS analysis

Flag-tagged GSDMD immunoprecipitates prepared from whole-cell lysates or gel-filtrated fractions were resolved on SDS-PAGE gels, and protein bands were excised. The samples were digested with trypsin, and then subject to LC-MS/MS Analysis. Swissprot_Human mass spectra were used as the standard reference. Trypsin/P was used for cleavage. MS data was captured and analyzed using Micrometer Biotech and Maxquant, respectively.

### Statistical analysis

The values are presented as mean ± SD. Data was analyzed using GraphPad Prism 8.0. Difference between experimental groups was assessed using Student’s *t*-test or one-way ANOVA. All experiments were performed independently at least three times. Statistical significance was set at *P* < 0.05, 0.01 or 0.001. \**P* represents *P*< 0.05, \*\**P* represents *P* < 0.01 and \*\*\**P* represents *P* < 0.001.

## Acknowledgments

This work was financially supported by the Zhejiang Provincial Key R&D Program of China (2021C02049), the National Natural Science Foundation of China (32072817) , the Scientific Research Fund of Zhejiang Provincial Education Department (Y202045613), the Zhejiang Provincial Natural Science Foundation of China (LY18C180001, LY21C180001), and the Intergovernmental Collaborative Project in S & T Innovation under the National Key R & D Program (Grant No. 2018YFE0111700).

We thank Dr. Ying Shan and Weiren Dong in the Shared Experimental Platform for Core Instruments, College of Animal Sciences, Zhejiang University for assistance with analysis of laser confocal microscopy imaging.

## Conflict of interest

The authors declare that they have no conflict of interest.

---

**Supplementary Figure 1 A,** Prediction of GSDMD ubiquitination by E3 ubiquitin ligase using the Ubibrowser software. **B,** MS/MS analysis for representative sequences of Human SYVN1 alleles.

**Supplementary Figure 2 A,** Bright-field and epifluorescent microscopy of HEK293T cells after PI staining. Scale: 1 bar represents 50 μm. **B,** HEK293T cells were transfected with Myc-SYVN1 and p3XFlag-hGSDMD-FL and thereafter for 18 h or 24 h with Caspase-1/4. **C,** HEK293T cell were transfected with plasmids as shown. PI analysis was performed after 24 h transfection. Scale: 1 bar represents 50 μm. **D-E,** HEK293T cells transfected with varied dose of hGSDMD-p30 and Ub. The supernatants were collected and analyzed after 24 h transfection for LDH release assay (D). HEK293T cells were observed under bright-field and epifluorescent microscopy after PI staining (E). Scale: 1 bar represents 100 μm.

**Supplementary Figure 3 ,** HEK293T cells under a fluorescence microscopy after PI staining. The cells were first transfected with siSYVN1 (50 nM) or corresponding control and thereafter with hGSDMD-p30 after 48 h. Scale: 1 bar represents 50 μm.

**Supplementary Figure 4 Putative sites on hGSDMD**. **A,** Representative MS/MS spectra of ubiquitinated hGSDMD peptides. Protein samples were recovered from the gel, enzymatically digested and analyzed using Mass Spectrometry (MS). **B,** Ubiquitination patterns of GSDMD. Red areas represent ubiquitination sites predicted by Ubibrowser software, whereas the green sites represent ubiquitination sites identified by MS.

**Supplementary Figure 5 A-B,** PI staining analysis of HEK293T cells after 24 h transfection with indicated plasmids. Scale: 1 bar represents 50 μm.

## Notes

### Competing Interest Statement

The authors have declared no competing interest.

## References

1. Shi J, Zhao Y, Wang Y, Gao W, Ding J, Li P, et al. Inflammatory caspases are innate immune receptors for intracellular LPS. Nature. 2014; 514(7521): 187–92.

2. Kayagaki N, Warming S, Lamkanfi M, Vande Walle L, Louie S, Dong J, et al. Non-canonical inflammasome activation targets caspase-11. Nature. 2011; 479(7371): 117–21.

3. Bergsbaken T, Fink SL, Cookson BT. Pyroptosis: host cell death and inflammation. Nat Rev Microbiol. 2009; 7(2): 99–109.

4. Broz P. Immunology: Caspase target drives pyroptosis. Nature. 2015; 526(7575): 642–643.

5. Shi J, Zhao Y, Wang K, Shi X, Wang Y, Huang H, et al. Cleavage of GSDMD by inflammatory caspases determines pyroptotic cell death. Nature. 2015; 526(7575): 660–5.

6. Kayagaki N, Stowe IB, Lee BL, O’Rourke K, Anderson K, Warming S, et al. Caspase-11 cleaves gasdermin D for non-canonical inflammasome signalling. Nature. 2015; 526(7575): 666–71.

7. Shi JJ, Zhao Y, Wang YP, Gao WQ, Ding JJ, Li P, et al. Inflammatory caspases are innate immune receptors for intracellular LPS. Nature. 2014; 514(7521): 187–92.

8. Rathinam VA, Vanaja SK, Fitzgerald KA. Regulation of inflammasome signaling. Nat Immunol. 2012; 13(4): 333–42.

9. Broz P, Dixit VM. Inflammasomes: mechanism of assembly, regulation and signalling. Nature Reviews Immunology. 2016; 16(7): 407–20.

10. Lamkanfi M, Dixit VM. Inflammasomes and Their Roles in Health and Disease. Annual Review of Cell and Developmental Biology, Vol 28. 2012; 28: 137–61.

11. de Vasconcelos NM, Lamkanfi M. Recent Insights on Inflammasomes, Gasdermin Pores, and Pyroptosis. Cold Spring Harb Perspect Biol. 2020; 12(5): a036392.

12. Yang Y, Wang H, Kouadir M, Song H, Shi F. Recent advances in the mechanisms of NLRP3 inflammasome activation and its inhibitors. Cell Death Dis. 2019; 10(2): 128.

13. Kayagaki N, Stowe IB, Lee BL, O’Rourke K, Anderson K, Warming S, et al. Caspase-11 cleaves gasdermin D for non-canonical inflammasome signalling. Nature. 2015; 526(7575): 666–71.

14. Micaroni M, Stanley AC, Khromykh T, Venturato J, Wong CX, Lim JP, et al. Rab6a/a’ are important Golgi regulators of pro-inflammatory TNF secretion in macrophages. PLoS One. 2013; 8(2): e57034.

15. Aglietti RA, Estevez A, Gupta A, Ramirez MG, Liu PS, Kayagaki N, et al. GsdmD p30 elicited by caspase-11 during pyroptosis forms pores in membranes. Proc Natl Acad Sci U S A. 2016; 113(28): 7858–63.

16. Ding J, Wang K, Liu W, She Y, Sun Q, Shi J, et al. Erratum: Pore-forming activity and structural autoinhibition of the gasdermin family. Nature. 2016; 540(7631): 150.

17. Ding J, Wang K, Liu W, She Y, Sun Q, Shi J, et al. Pore-forming activity and structural autoinhibition of the gasdermin family. Nature. 2016; 535(7610): 111–6.

18. Liu X, Zhang Z, Ruan J, Pan Y, Magupalli VG, Wu H, et al. Inflammasome-activated gasdermin D causes pyroptosis by forming membrane pores. Nature. 2016; 535(7610): 153–8.

19. Sborgi L, Ruhl S, Mulvihill E, Pipercevic J, Heilig R, Stahlberg H, et al. GSDMD membrane pore formation constitutes the mechanism of pyroptotic cell death. EMBO J. 2016; 35(16): 1766–78.

20. Cockram PE, Kist M, Prakash S, Chen SH, Wertz IE, Vucic D. Ubiquitination in the regulation of inflammatory cell death and cancer. Cell Death Differ. 2021; 28(2): 591–605.

21. Bednash JS, Mallampalli RK. Regulation of inflammasomes by ubiquitination. Cell Mol Immunol. 2016; 13(6): 722–8.

22. Paik S, Kim JK, Silwal P, Sasakawa C, Jo EK. An update on the regulatory mechanisms of NLRP3 inflammasome activation. Cell Mol Immunol. 2021; 18(5): 1141–60.

23. Song H, Liu B, Huai W, Yu Z, Wang W, Zhao J, et al. The E3 ubiquitin ligase TRIM31 attenuates NLRP3 inflammasome activation by promoting proteasomal degradation of NLRP3. Nat Commun. 2016; 7: 13727.

24. Guan K, Wei CW, Zheng ZR, Song T, Wu FX, Zhang YH, et al. MAVS Promotes Inflammasome Activation by Targeting ASC for K63-Linked Ubiquitination via the E3 Ligase TRAF3. Journal of Immunology. 2015; 194(10): 4880–90.

25. Liu QJ, Zhang SH, Sun ZJ, Guo X, Zhou H. E3 ubiquitin ligase Nedd4 is a key negative regulator for non-canonical inflammasome activation. Cell Death and Differentiation. 2019; 26(11): 2386–99.

26. Zhang L, Liu Y, Wang B, Xu G, Yang Z, Tang M, et al. POH1 deubiquitinates pro-interleukin-1beta and restricts inflammasome activity. Nat Commun. 2018; 9(1): 4225.

27. Yan YQ, Jiang W, Liu L, Wang XQ, Ding C, Tian ZG, et al. Dopamine Controls Systemic Inflammation through Inhibition of NLRP3 Inflammasome. Cell. 2015; 160(1-2): 62–73.

28. Humphries F, Bergin R, Jackson R, Delagic N, Wang BW, Yang S, et al. The E3 ubiquitin ligase Pellino2 mediates priming of the NLRP3 inflammasome. Nature Communications. 2018; 9(1):1560

29. Guo Y, Li LJ, Xu T, Guo XM, Wang CM, Li YH, et al. HUWE1 mediates inflammasome activation and promotes host defense against bacterial infection. Journal of Clinical Investigation. 2020; 130(12): 6301–16.

30. Hansen JM, de Jong MF, Wu Q, Zhang LS, Heisler DB, Alto LT, et al. Pathogenic ubiquitination of GSDMB inhibits NK cell bactericidal functions. Cell. 2021; 184(12): 3178–3191 e3118.

31. Xia S. Biological mechanisms and therapeutic relevance of the gasdermin family. Mol Aspects Med. 2020; 76: 100890.

32. Hasegawa D, Fujii R, Yagishita N, Matsumoto N, Aratani S, Izumi T, et al. E3 Ubiquitin Ligase Synoviolin Is Involved in Liver Fibrogenesis. Plos One. 2010; 5(10) e13590.

33. Gao B, Lee SM, Chen A, Zhang J, Zhang DD, Kannan K, et al. Synoviolin promotes IRE1 ubiquitination and degradation in synovial fibroblasts from mice with collagen-induced arthritis. EMBO Rep. 2008; 9(5): 480–5.

34. Yang H, Qiu Q, Gao B, Kong S, Lin Z, Fang D. Hrd1-mediated BLIMP-1 ubiquitination promotes dendritic cell MHCII expression for CD4 T cell priming during inflammation. J Exp Med. 2014; 211(12): 2467–79.

35. Luo Z, Liu LF, Jiang YN, Tang LP, Li W, Ouyang SH, et al. Novel insights into stress-induced susceptibility to influenza: corticosterone impacts interferon-beta responses by Mfn2-mediated ubiquitin degradation of MAVS. Signal Transduction and Targeted Therapy. 2020; 5(1): 202.

36. Xu Y, Zhao F, Qiu Q, Chen K, Wei J, Kong Q, et al. The ER membrane-anchored ubiquitin ligase Hrd1 is a positive regulator of T-cell immunity. Nat Commun. 2016; 7: 12073.

37. Kong S, Yang Y, Xu Y, Wang Y, Zhang Y, Melo-Cardenas J, et al. Endoplasmic reticulum-resident E3 ubiquitin ligase Hrd1 controls B-cell immunity through degradation of the death receptor CD95/Fas. Proc Natl Acad Sci U S A. 2016; 113(37): 10394–9.

38. Lu Y, Qiu Y, Chen P, Chang H, Guo L, Zhang F, et al. ER-localized Hrd1 ubiquitinates and inactivates Usp15 to promote TLR4-induced inflammation during bacterial infection. Nat Microbiol. 2019; 4(12): 2331–46.

39. Sollberger G, Choidas A, Burn GL, Habenberger P, Di Lucrezia R, Kordes S, et al. Gasdermin D plays a vital role in the generation of neutrophil extracellular traps. Sci Immunol. 2018; 3(26): eaar6689.

40. Hu JJ, Liu X, Xia S, Zhang Z, Zhang Y, Zhao J, et al. FDA-approved disulfiram inhibits pyroptosis by blocking gasdermin D pore formation. Nat Immunol. 2020; 21(7): 736–745.

41. Lei X, Zhang Z, Xiao X, Qi J, He B, Wang J. Enterovirus 71 Inhibits Pyroptosis through Cleavage of Gasdermin D. J Virol. 2017; 91(18): e01069–17.

42. Kaneko M, Ishiguro M, Niinuma Y, Uesugi M, Nomura Y. Human HRD1 protects against ER stress-induced apoptosis through ER-associated degradation. FEBS Lett. 2002; 532(1-2): 147–52.

43. van de Weijer ML, Bassik MC, Luteijn RD, Voorburg CM, Lohuis MAM, Kremmer E, et al. A high-coverage shRNA screen identifies TMEM129 as an E3 ligase involved in ER-associated protein degradation. Nature Communications. 2014; 5: 3832.

44. Swatek KN, Komander D. Ubiquitin modifications. Cell Res. 2016; 26(4): 399–422.

45. Yau RG, Doerner K, Castellanos ER, Haakonsen DL, Werner A, Wang N, et al. Assembly and Function of Heterotypic Ubiquitin Chains in Cell-Cycle and Protein Quality Control. Cell. 2017; 171(4): 918–933 e920.

46. Oh E, Akopian D, Rape M. Principles of Ubiquitin-Dependent Signaling. Annu Rev Cell Dev Biol. 2018; 34: 137–62.

47. Deshaies RJ, Joazeiro CA. RING domain E3 ubiquitin ligases. Annu Rev Biochem. 2009; 78: 399–434.

48. Wang L, Wu JH, Li J, Yang H, Tang TQ, Liang HJ, et al. Host-mediated ubiquitination of a mycobacterial protein suppresses immunity. Nature. 2020; 577(7792): 682–88.

49. Radivojac P, Vacic V, Haynes C, Cocklin RR, Mohan A, Heyen JW, et al. Identification, analysis, and prediction of protein ubiquitination sites. Proteins. 2010; 78(2): 365–80.

50. Zheng Y, Liu Q, Wu Y, Ma L, Zhang Z, Liu T, et al. Zika virus elicits inflammation to evade antiviral response by cleaving cGAS via NS1-caspase-1 axis. EMBO J. 2018; 37(18).

51. Tang J, Tu S, Lin G, Guo H, Yan C, Liu Q, et al. Sequential ubiquitination of NLRP3 by RNF125 and Cbl-b limits inflammasome activation and endotoxemia. J Exp Med. 2020; 217(4): e20182091.

52. Rathkey JK, Benson BL, Chirieleison SM, Yang J, Xiao TS, Dubyak GR, et al. Live-cell visualization of gasdermin D-driven pyroptotic cell death. Journal of Biological Chemistry. 2017; 292(35): 14649–58.

53. Orning P, Lien E, Fitzgerald KA. Gasdermins and their role in immunity and inflammation. Journal of Experimental Medicine. 2019; 216(11): 2453–65.

54. Kayagaki N, Warming S, Lamkanfi M, Vande Walle L, Louie S, Dong J, et al. Non-canonical inflammasome activation targets caspase-11. Nature. 2011; 479(7371): 117–U146.

55. Liu Q, Zhang S, Sun Z, Guo X, Zhou H. E3 ubiquitin ligase Nedd4 is a key negative regulator for non-canonical inflammasome activation. Cell Death Differ. 2019; 26(11): 2386–99.

56. Palazon-Riquelme P, Worboys JD, Green J, Valera A, Martin-Sanchez F, Pellegrini C, et al. USP7 and USP47 deubiquitinases regulate NLRP3 inflammasome activation. EMBO Rep. 2018; 19(10): e44766.

57. Gao B, Calhoun K, Fang D. The proinflammatory cytokines IL-1beta and TNF-alpha induce the expression of Synoviolin, an E3 ubiquitin ligase, in mouse synovial fibroblasts via the Erk1/2-ETS1 pathway. Arthritis Res Ther. 2006; 8(6): R172.

58. Rathkey JK, Xiao TS, Abbott DW. Human polymorphisms in GSDMD alter the inflammatory response. J Biol Chem. 2020; 295(10): 3228–38.

59. Burr ML, van den Boomen DJH, Bye H, Antrobus R, Wiertz EJ, Lehner PJ. MHC class I molecules are preferentially ubiquitinated on endoplasmic reticulum luminal residues during HRD1 ubiquitin E3 ligase-mediated dislocation. Proceedings of the National Academy of Sciences of the United States of America. 2013; 110(35): 14290–5.

60. Shi Y, Wang H, Zheng M, Xu W, Yang Y, Shi F. Ginsenoside Rg3 suppresses the NLRP3 inflammasome activation through inhibition of its assembly. FASEB J. 2020; 34(1): 208–21.

