## Supplemental Figure1 for "E3 ubiquitin ligase SYVN1 is a key positive regulator for GSDMD-mediated pyroptosis"

A

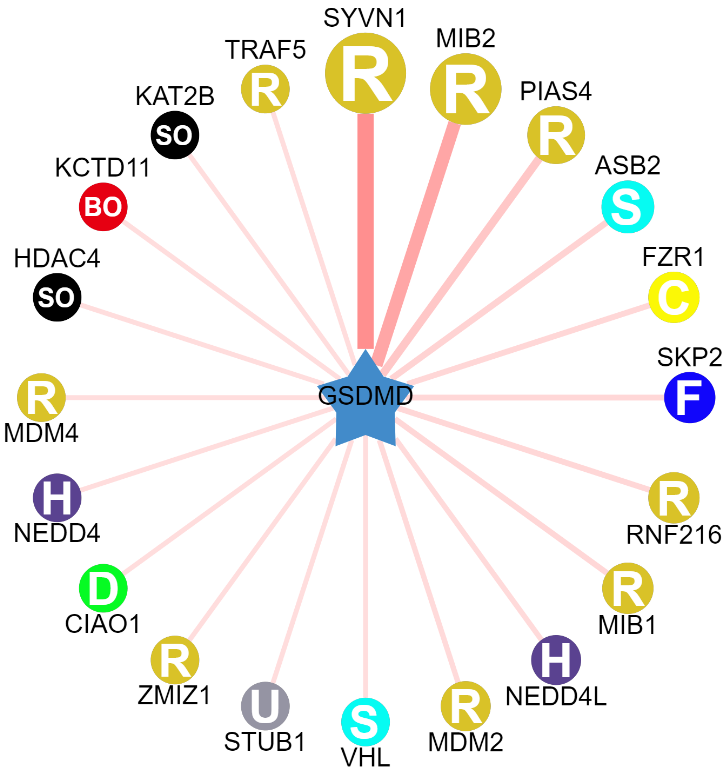


B

**Table 1 | Representative peptide sequences corresponding to Human SYVN1, detected in MS/MS analysis**

| No. | Sequence of peptide | Start position | End position |
|---|---|---|---|
| 1 | HQFYPTVVYLTK | 27 | 38 |
| 2 | AAEMEHLLER | 71 | 80 |
| 3 | SPNISWLFHCR | 130 | 140 |
| 4 | VHTFPLFAIR | 237 | 246 |
| 5 | KAVTDAIMSR | 257 | 266 |
| 6 | EEMVTGAK | 296 | 303 |
| 7 | RLPCNHIFHTSCLR | 304 | 317 |
| 8 | QQTCPTCR | 323 | 330 |
