## Supplementary figures and images for "E3 ubiquitin ligase SYVN1 is a key positive regulator for GSDMD-mediated pyroptosis"

### Supplemental Figure2

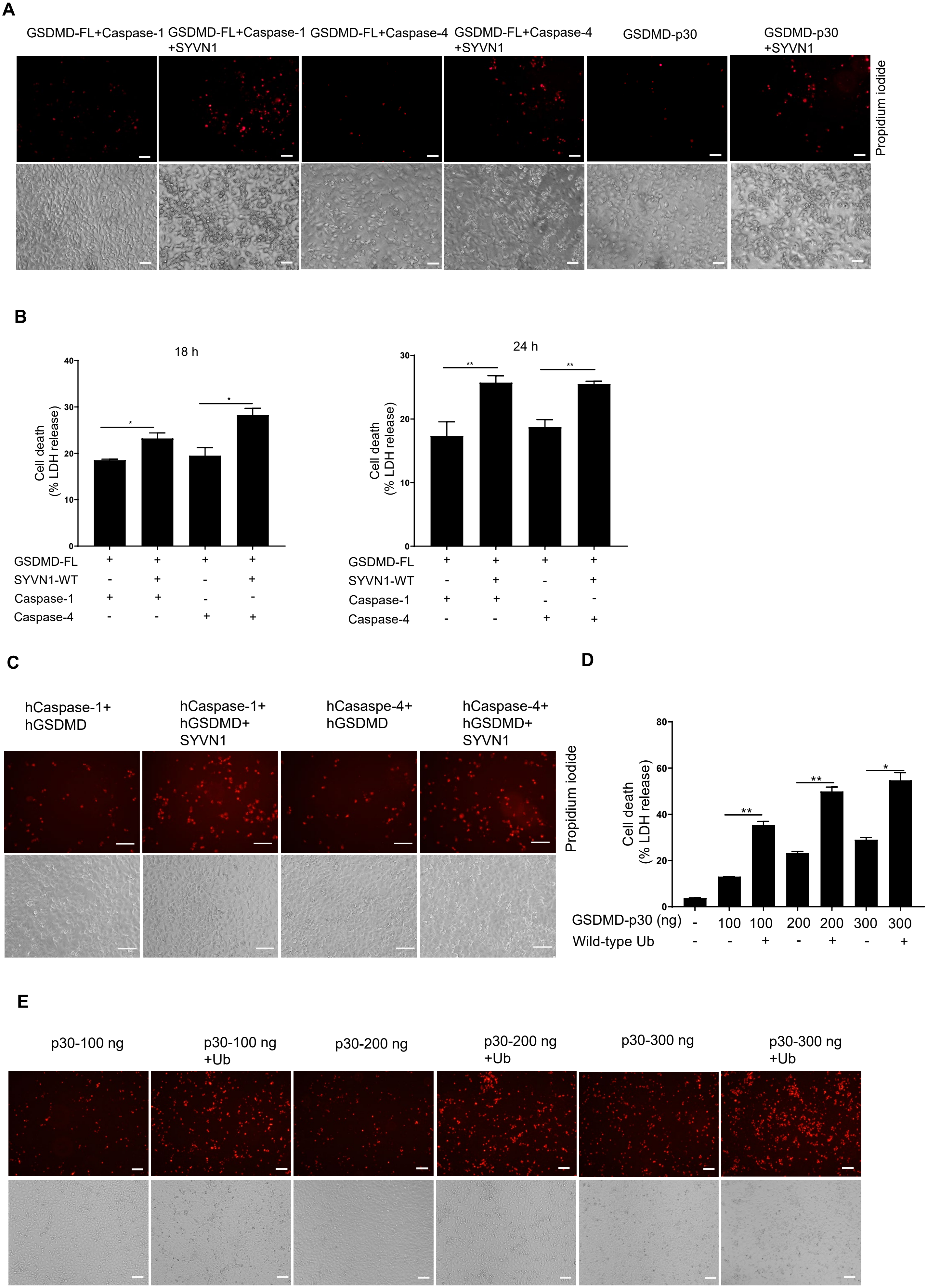

### Supplemental Figure3

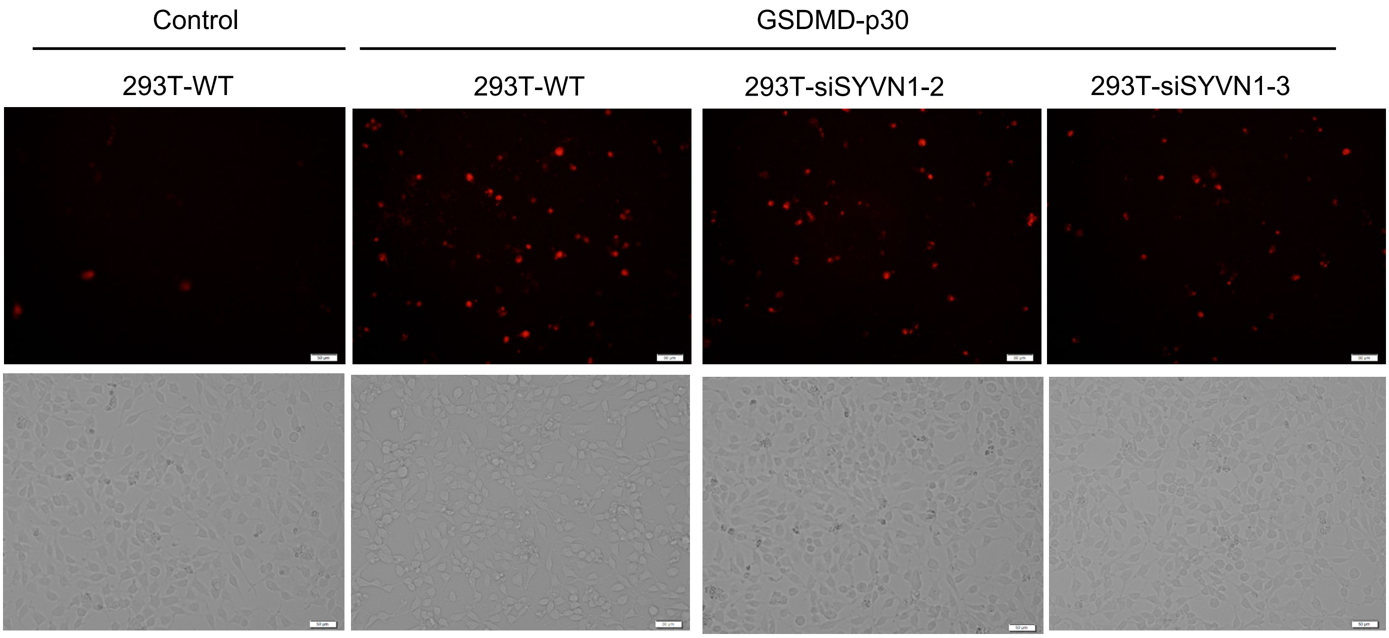

### Supplemental Figure4

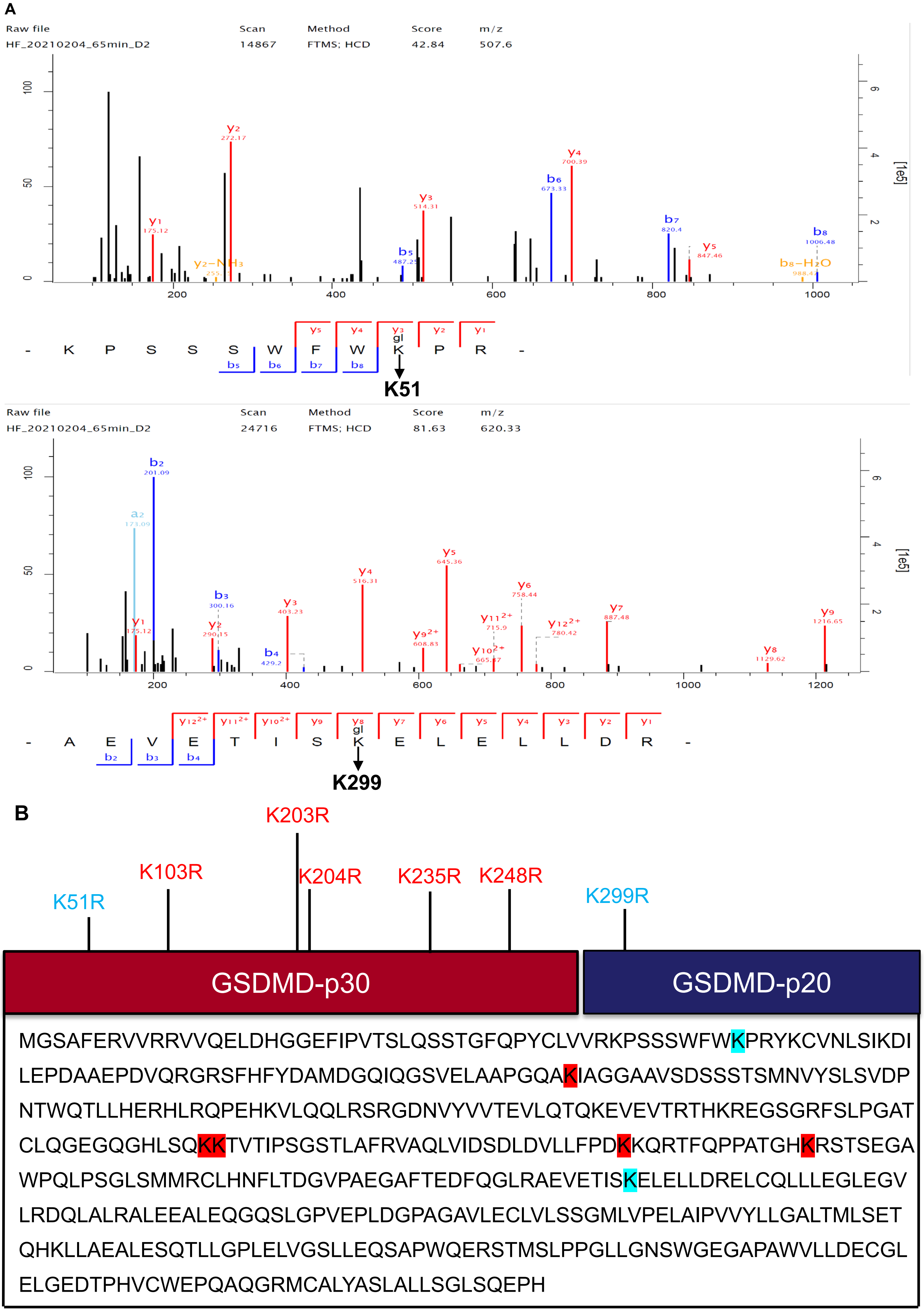

### Supplemental Figure5

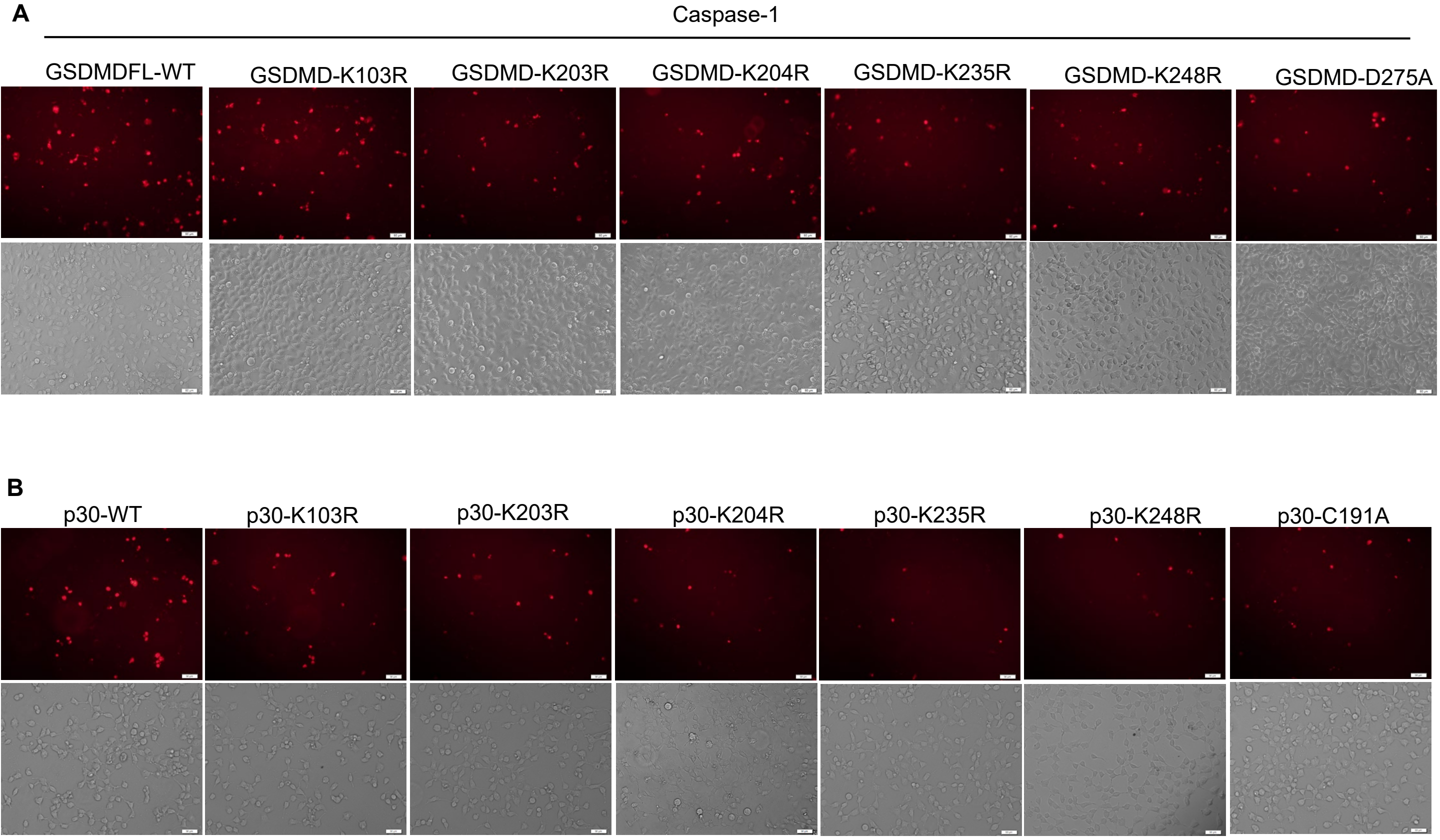
