## Supplemental Table1 for "E3 ubiquitin ligase SYVN1 is a key positive regulator for GSDMD-mediated pyroptosis"

**Table 1** Primers used in this study for plasmids construction.

| **Primers** | **Sequences** |
| --- | --- |
| p3XFlag-hGSDMD-FL forward | 5’-AAGGATGACGATGACAAGCTTATGGGGTCGGCCTTTGAG-3’ |
| p3XFlag-hGSDMD-FL reverse | 5’-ATCAGATCTATCGATGAATTCCTAGTGGGGCTCCTGGCTCA-3’ |
| pCMV-Myc-hCaspase-1 forward | 5’-GGAGGCCCGAATTCGGTCGACCATGGCCGACAAGGTCCTG-3’ |
| pCMV-Myc-hCaspase-1 reverse | 5’-CATGTCTGGATCCCCGCGGCCGCTTAATGTCCTGGGAAGAGGTAGAAA-3’ |
| pCMV-Myc-hCaspase-4 forward | 5’-GGAGGCCCGAATTCGGTCGACCATGGCAGAAGGCAACCACA-3’ |
| pCMV-Myc-hCaspase-4 reverse | 5’-CATGTCTGGATCCCCGCGGCCGCTCAATTGCCAGGAAAGAGGTAGA-3’ |
| p3XFlag-h Caspase-1 forward | 5’-CAAGCTTGCGGCCGCGAATTCAGCCACCATGGCCGACAA-3’ |
| p3XFlag-hCaspase-1 reverse | 5’-CCTCTAGAGTCGACTGGTACCTTAATGTCCTGGGAAGAGGTAGAAA-3’ |
| p3XFlag- hCaspase-4 forward | 5’-AAGGATGACGATGACAAGCTTATGGCAGAAGGCAACCACAG-3’ |
| p3XFlag- hCaspase-4 reverse | 5’-ATCAGATCTATCGATGAATTCTCAATTGCCAGGAAAGAGGTAGA-3’ |
| pcDNA3.1-hGSDMD-p30-Myc forward | 5’-CCAAGCTGGCTAGTTAAGCTTATGGGGTCGGCCTTTGAG-3’ |
| pcDNA3.1-hGSDMD-p30-Myc reverse | 5’-GAGTTTTTGTTCGAAGGGCCCATCTGTCAGGAAGTTGTGGAGGC-3’ |
| p3×Flag-N-pGSDMD-FL forward | 5’-CAAGCTTGCGGCCGCGAATTCTATGGCATCAGCCTTTGAGAGG-3’ |
| p3×Flag-N-pGSDMD-FL reverse | 5’-TGCCACCCGGGATCCTCTAGACTAGCAGAGCTGGCTGAGCC-3’ |
| p3×Flag-N-mGSDMD-FL forward | 5’-AAGGATGACGATGACAAGCTTATGCCATCGGCCTTTGAGA-3’ |
| p3×Flag-N-mGSDMD-FL reverse | 5’-ATCAGATCTATCGATGAATTCCTAACAAGGTTTCTGGCCTAGACTT-3’ |
| p3XFlag-N-SYVN1 forward | 5’-AAGGATGACGATGACAAGCTTATGTTCCGCACGGCAGTG-3’ |
| p3XFlag-N-SYVN1 reverse | 5’-ATCAGATCTATCGATGAATTCTCAGTGGGCAACAGGAGACTCC-3’ |
| p3XFlag-N-SYVN1-C329S forward | 5’-ACCAGCCGTATGGATGTCCTTCGTGCATCGCT-3’ |
| p3XFlag-N-SYVN1-C329S reverse | 5’-ACATCCATACGGCTGGTGGGGCAGGTCTGCTG-3’ |
| pcDNA3.1-C-SYVN1-Myc forward | 5’-CTTGGTACCGAGCTCGGATCCGCCACCATGTTCCGCACG-3’ |
| pcDNA3.1-C-SYVN1-Myc reverse | 5’-GAGTTTTTGTTCGAAGGGCCCGTGGGCAACAGGAGACTCCA-3’ |
| p3XFlag-hGSDMD-FL-K51R-forward | 5’-TTCTGGAGACCCCGTTATAAGTGTGTCAACCT -3’ |
| p3XFlag-hGSDMD-FL-K51R reverse | 5’-TAACGGGGTCTCCAGAACCATGAGCTTGAGGG-3’ |
| p3XFlag-hGSDMD-FL-K103R forward | 5’-AGCCCCAGGACAGGCAAGGATCGCAGGCGGGGCCG-3’ |
| p3XFlag-hGSDMD-FL-K103R reverse | 5’-CGGCCCCGCCTGCGATCCTTGCCTGTCCTGGGGCT-3’ |
| p3XFlag-hGSDMD-FL-K203R forward | 5’-AGGGCCAGGGCCATCTGAGCCAGAGGAAGACGGTCACCATCCCCTCAGG -3’ |
| p3XFlag-hGSDMD-FL-K203R reverse | 5’-CCTGAGGGGATGGTGACCGTCTTCCTCTGGCTCAGATGGCCCTGGCCCT -3’ |
| p3XFlag-hGSDMD-FL-K204R forward | 5’-GCCAGGGCCATCTGAGCCAGAAGAGGACGGTCACCATCCCCTCAGGCAG -3’ |
| p3XFlag-hGSDMD-FL-K204R reverse | 5’-CTGCCTGAGGGGATGGTGACCGTCCTCTTCTGGCTCAGATGGCCCTGGC -3’ |
| p3XFlag-hGSDMD-FL-K235R forward | 5’-TGGACGTCCTTCTCTTCCCGGATAGGAAGCAGAGGACCTTCCAGCCACC -3’ |
| p3XFlag-hGSDMD-FL-K235R reverse | 5’-GGTGGCTGGAAGGTCCTCTGCTTCCTATCCGGGAAGAGAAGGACGTCCA -3’ |
| p3XFlag-hGSDMD-FL-K248R forward | 5’-CCACCCGCGACAGGCCACAGGCGTTCCACGAGCGAAGGC-3’ |
| p3XFlag-hGSDMD-FL-K248R reverse | 5’-GCCTTCGCTCGTGGAACGCCTGTGGCCTGTCGCGGGTGG-3’ |
| p3XFlag-hGSDMD-FL-K299R forward | 5’-ATCTCCAGGGAACTGGAGCTTTTGGACAGAGA-3’ |
| p3XFlag-hGSDMD-FL-K299R reverse | 5’-TCCAGTTCCCTGGAGATGGTCTCCACCTCTGC-3’ |
| p3XFlag-hGSDMD-FL-D275A forward | 5’-GCCTCCACAACTTCCTGACAGCTGGGGTCCCTGCGGAGGGGGC-3’ |
| p3XFlag-hGSDMD-FL- D275A reverse | 5’-GCCCCCTCCGCAGGGACCCCAGCTGTCAGGAAGTTGTGGAGGC-3’ |
| pCDNA3.1-Myc-C-hGSDMD-p30-C191A forward | 5’-CACGGCCTTGCAGGGTGAGGGCCAGGGCCATCT-3’ |
| pCDNA3.1-Myc-C-hGSDMD-p30-C191A reverse | 5’-TCACCCTGCAAGGCCGTGGCTCCGGGCAGGGAA-3’ |
