## Supplemental Table2 for "E3 ubiquitin ligase SYVN1 is a key positive regulator for GSDMD-mediated pyroptosis"

**Table 2** siRNA sequence for the SYVN1 oligonucelotide.

| **Name** | **Sequences** |
| --- | --- |
| *siSYVN#1* forward | 5’-GCCGCAUUGUCUCUCUUAUTT-3’ |
| *siSYVN#1* reverse | 5’-AUAAGAGAGACAAUGCGGCTT-3’ |
| *siSYVN#2* forward | 5’-GUGCUCACCAUCUUCAUCATT-3’ |
| *siSYVN#2* reverse | 5’-UGAUGAAGAUGGUGAGCACTT-3’ |
| *siSYVN#3* forward | 5’-CAGGCUUCAUCAAGGUUCUTT-3’ |
| *siSYVN#3* reverse | 5’-AGAACCUUGAUGAAGCCUGTT-3’ |
| Negative control forward | 5’-UUCUCCGAACGUGUCACGUTT-3 |
| Negative control reverse | 5’-ACGUGACACGUUCGGAGAATT-3’ |
| *GAPDH* forward | 5’-UGACCUCAACUACAUGGUUTT-3’ |
| *GAPDH* reverse | 5’-AACCAUGUAGUUGAGGUCATT-3’ |
